# Zinc binding inhibits cellular uptake and antifungal activity of Histatin-5 in *Candida albicans*

**DOI:** 10.1101/2022.06.11.494584

**Authors:** Joanna X. Campbell, Sean Gao, Keerthi S. Anand, Katherine J. Franz

## Abstract

Histatin-5 (Hist-5) is a polycationic, histidine-rich antimicrobial peptide with potent antifungal activity against the opportunistic fungal pathogen *Candida albicans*. Hist-5 has the ability to bind metals in vitro and metals have been shown to alter the fungicidal activity of the peptide. Previous reports on the effect of Zn^2+^ on Hist-5 activity have been varied and seemingly contradictory. Here we present data elucidating the dynamic role Zn^2+^ plays as an inhibitory switch to regulate Hist-5 fungicidal activity. A novel fluorescently labeled Hist-5 peptide (Hist-5*) was developed to visualize changes in internalization and localization of the peptide as a function of metal availability in the growth medium. Hist-5* was verified for use as a model peptide and retained antifungal activity and mode of action similar to native Hist-5. Cellular growth assays showed that Zn^2+^ had a concentration-dependent inhibitory effect on Hist-5 antifungal activity. Imaging by confocal microscopy revealed that equimolar concentrations of Zn^2+^ kept the peptide localized along the cell periphery rather than internalizing, thus preventing cytotoxicity and membrane disruption. However, the Zn-induced decrease in Hist-5 activity and uptake was rescued by decreasing Zn^2+^ availability upon addition of a metal chelator EDTA or S100A12, a Zn-binding protein involved in the innate immune response. These results lead us to suggest a model wherein commensal *C. albicans* may exist in harmony with Hist-5 at concentrations of Zn^2+^ that inhibit peptide internalization and antifungal activity. Activation of host immune processes that initiate Zn-sequestering mechanisms of nutritional immunity could trigger Hist-5 internalization and cell killing.

## Introduction

Histatin-5 (Hist-5) is a histidine-rich peptide that is naturally produced in the salivary glands of higher primates as a part of the innate immune system.^1^ Antimicrobial activity of Hist-5 has been reported against a variety of bacterial and fungal species, including the opportunistic fungal pathogen, *Candida albicans* (*C. albicans*).^1–5^ Fungal heat shock proteins Ssa1 and Ssa2 have been identified as cell-surface receptors for Hist-5, with subsequent intracellular translocation utilizing polyamine transporters Dur3 and 31.^6–10^ Once internalized, Hist-5 treatment ultimately leads to cell cycle arrest, volume dysregulation, formation of reactive oxygen species, and nonlytic efflux of ATP and other cytosolic small molecules and ions.^11–16^ While much is known about the antifungal outcomes of Hist-5 on *C. albicans*, the mechanisms of Hist-5 activity are not fully established. Of particular interest to us is how metal ions potentiate the mode of action of Hist-5. The amino acid sequence of Hist-5 (DSHAKRHHGYKRKFHEKHHSHRGY) possesses several metal-binding motifs capable of binding ions of multiple oxidation states of copper (Cu^+/2+^), iron (Fe^2+/3+^), and zinc (Zn^2+^) with varying affinities.^17–21^ We have shown previously that co-administration of Cu^2+^ salts with Hist-5 improved its candidacidal activity, while addition of Fe^3+^ decreased its activity.^19, 20^ Reports on the effects of Zn^2+^ on the activity of Hist-5, however, have been varied and seemingly contradictory.

Hist-5 has been reported to bind up to two equivalents of Zn^2+^, one of which is putatively at the HEXXH site, a prominent zinc-binding motif found in larger metalloproteins.^17, 18, 20–23^ Zn^2+^ binding has been reported to induce conformational changes in Hist-5, favoring alpha-helical secondary structure and promoting dimerization and aggregation under some conditions.^24–27^ While some studies suggest that Zn^2+^ has little to no effect on Hist-5 activity,^20^ others claim Zn^2+^ causes a decrease in the antifungal activity of the Hist-5 derivative, P113.^28^ A recent study by Norris *et al.* reported an increase in Hist-5 antifungal activity at a 2:1 ratio of peptide to Zn^2+^.^25^ The authors subsequently followed up this study by showing that Hist-5+Zn^2+^ treatment not only promotes fungicidal activity but also has a role in maintaining commensalism of *C. albicans* in the oral cavity.^29^ While these studies suggest a role for Zn^2+^ in modulating the effects of Hist-5 on *C. albicans*, the overall response remains unclear and contradictory.^20, 25, 28^

Concentrations of Hist-5 and Zn^2+^ in saliva are dynamic ranging from 0.7 – 30 µM for Hist-5^30, 31^ and 0.0001 – 155 µM for Zn^2+^.^32–34^ These levels are subject to change based on a number of factors including age, diet, and health of the individual.^31, 33, 35–38^ Probing how Hist-5 operates across a dynamic range of Zn^2+^ availability may be important for gaining a comprehensive understanding of its effect in modulating Hist-5 activity. In this study, we therefore set out to evaluate the activity and localization of Hist-5 against *C. albicans* in liquid culture across a range of physiologically relevant peptide and Zn^2+^ concentrations. Our results demonstrate that increasing Zn^2+^ supplementation negatively affects Hist-5 antifungal activity. By using a novel fluorescent Hist-5 analogue, Hist-5*, we show that increasing the Zn^2+^ concentration promotes Hist-5 surface adhesion but inhibits peptide uptake into the cytosol. Furthermore, modulation of Zn^2+^ availability by extracellular metal-binding molecules reverses this Zn^2+^ inhibitory effect to recover Hist-5 cellular uptake and membrane disruption. These findings lead to a model in which the availability of Zn^2+^ may regulate the biological activity of Hist-5.

## Results

### Design and characterization of a fluorescent Hist-5 analogue

Metal availability and metal binding have a profound impact on Hist-5 structure and activity.^39, 40^ making retention of those properties crucial when developing modified Hist-5 peptides for study. Previous studies have utilized fluorescein or rhodamine to label Hist-5 to study peptide uptake and intracellular targets.^41, 42^ However, these labeling strategies conjugate the fluorophores at the N-terminus of the peptide, which would disrupt one of the recognized metal-binding sites of Hist-5, specifically the amino-terminal Cu^2+^, Ni^2+^ binding site. For our peptide design of Hist-5*, we opted for lower molecular weight fluorophores incorporated along the Hist- 5 sequence without disrupting metal-binding functionality. To minimize perturbations from bulky fluorophores, we chose to substitute the two tyrosine residues Y10 and Y24 with methoxycoumarin (Mca) and sulfamoylbenzofurazan (ABD) fluorophores, respectively (Figure 1A). These fluorescent amino acids were incorporated into the peptide sequence during solid-phase peptide synthesis, either directly as an fmoc-protected amino acid in the case of Mca, or via reaction with a unique cysteine in the case of ABD (Figure S1). A doubly-labeled version of Hist- 5 was chosen to enable potentially differential fluorescent response depending on metal type and binding site. Preliminary studies showed that tyrosine fluorescence specifically from Y10 was quenched by Cu^2+^, so this position was chosen for Mca installation. The ABD fluorophore was installed at the C-terminus to serve as a potential ratiometric handle.

**Figure 1.**
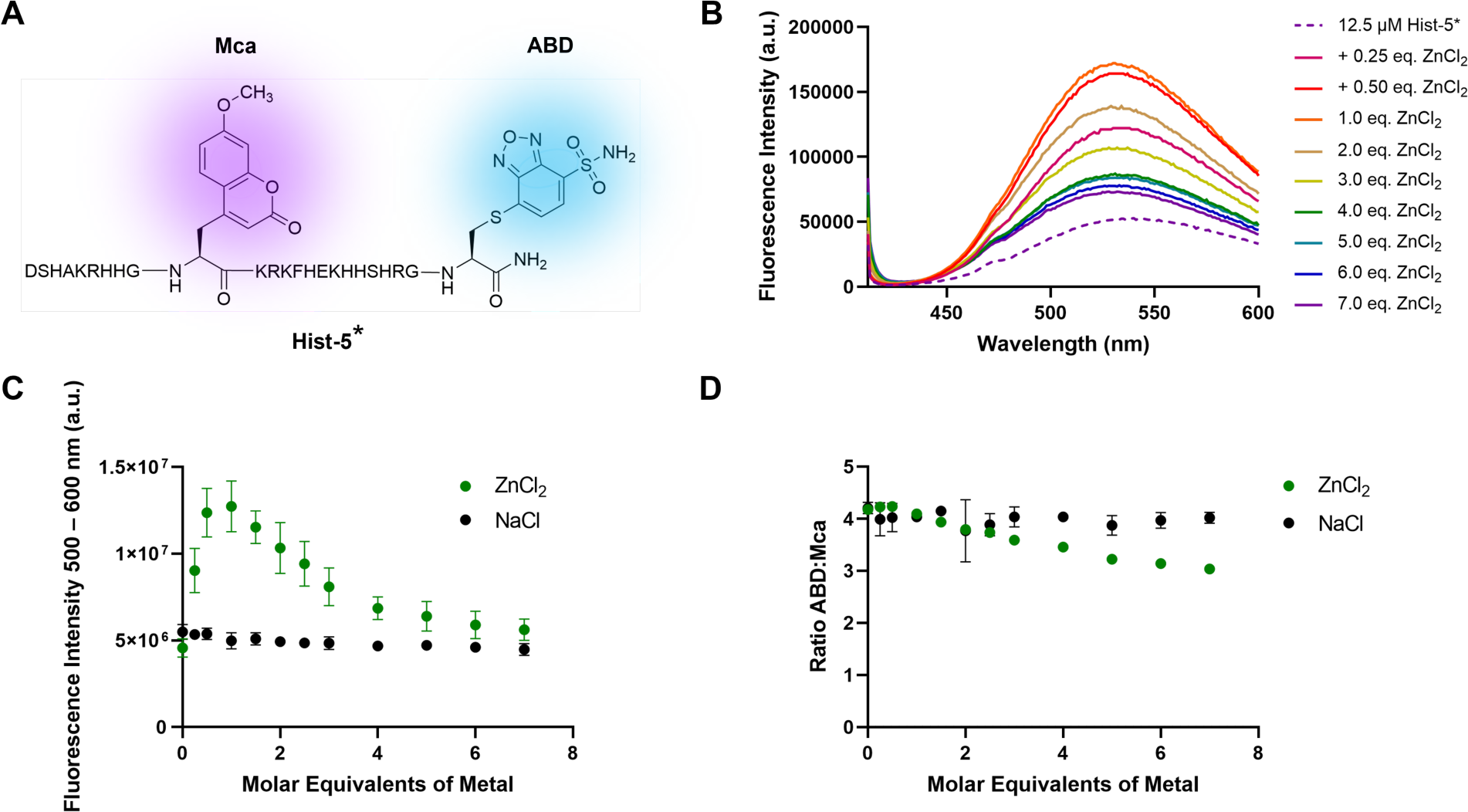
Sequence and characterization of Hist-5*. **(A)** Sequence of Hist-5* wherein Y10 and Y24 were replaced with methoxycoumarin (Mca, λ_ex_= 325 nm, λ_em_ = 400 nm) and sulfamoylbenzofurazan (ABD λ_em_ = 385 nm, λ_em_ = 510 nm) fluorophores. **(B)** Emission spectrum of Hist-5* excited at 405 nm upon titration of ZnCl_2_ into 12.5 µM Hist-5* in 1 mM potassium phosphate buffer (PPB) pH 7.4. **(C)** Fluorescence emission of the ABD fluorophore plotted as a function of added ZnCl_2_ (green) or NaCl (black). **(D)** Ratio of the fluorescence intensities of ABD to Mca as a function of added ZnCl_2_ (green) or NaCl (black).

To determine whether Hist-5* could be used to detect metal-dependent changes in fluorescence we probed the fluorescence response of the two fluorophores to Zn^2+^. In these experiments, Zn^2+^ was titrated into a solution of Hist-5* in 1 mM potassium phosphate buffer (PPB) pH 7.4 and the fluorescence emission from each fluorophore was monitored over two wavelength ranges, 412 – 499 and 500 – 600 nm, for Mca and ABD respectively. We chose to excite Hist-5* at 405 nm rather than at each fluorophore’s maximal excitation wavelength to mimic the excitation and emission parameters to be used in subsequent microscopy studies. As shown in Figures 1B and C, the majority of the fluorescence response from Hist-5* under these conditions can be attributed to the C-terminal ABD fluorophore. We observed an increase in Hist-5* fluorescence that peaked at 1 molar equivalent of added Zn^2+^, with subsequent additions of Zn^2+^ returning the emission intensity back to the original Hist-5* fluorescence. Titration with NaCl did not greatly affect Hist-5* fluorescence (Figures 1C and D), indicating that the increase in Hist-5* fluorescence was due to changes in Zn^2+^, not chloride. By plotting the ratio of the fluorescence intensities, ABD:Mca, we were able to detect metal-dependent changes to Hist-5* fluorescence (Figure 1D).

### Hist-5* retains antifungal activity and uptake similar to native Hist-5

The antifungal activity of Hist-5* against *C. albicans* was evaluated via a microdilution 96-well plate assay to determine the minimum inhibitory concentration (MIC) of the labeled peptide compared to unlabeled Hist-5. In these experiments, MIC is defined as the lowest concentration at which cell growth was no longer detected by optical density at 600 nm (OD_600_). We observed that cells treated with either Hist-5* or Hist-5 had an MIC of 25 µM (Figure 2A), indicating that the addition of the two fluorophores did not affect the antifungal activity of the peptide.

**Figure 2.**
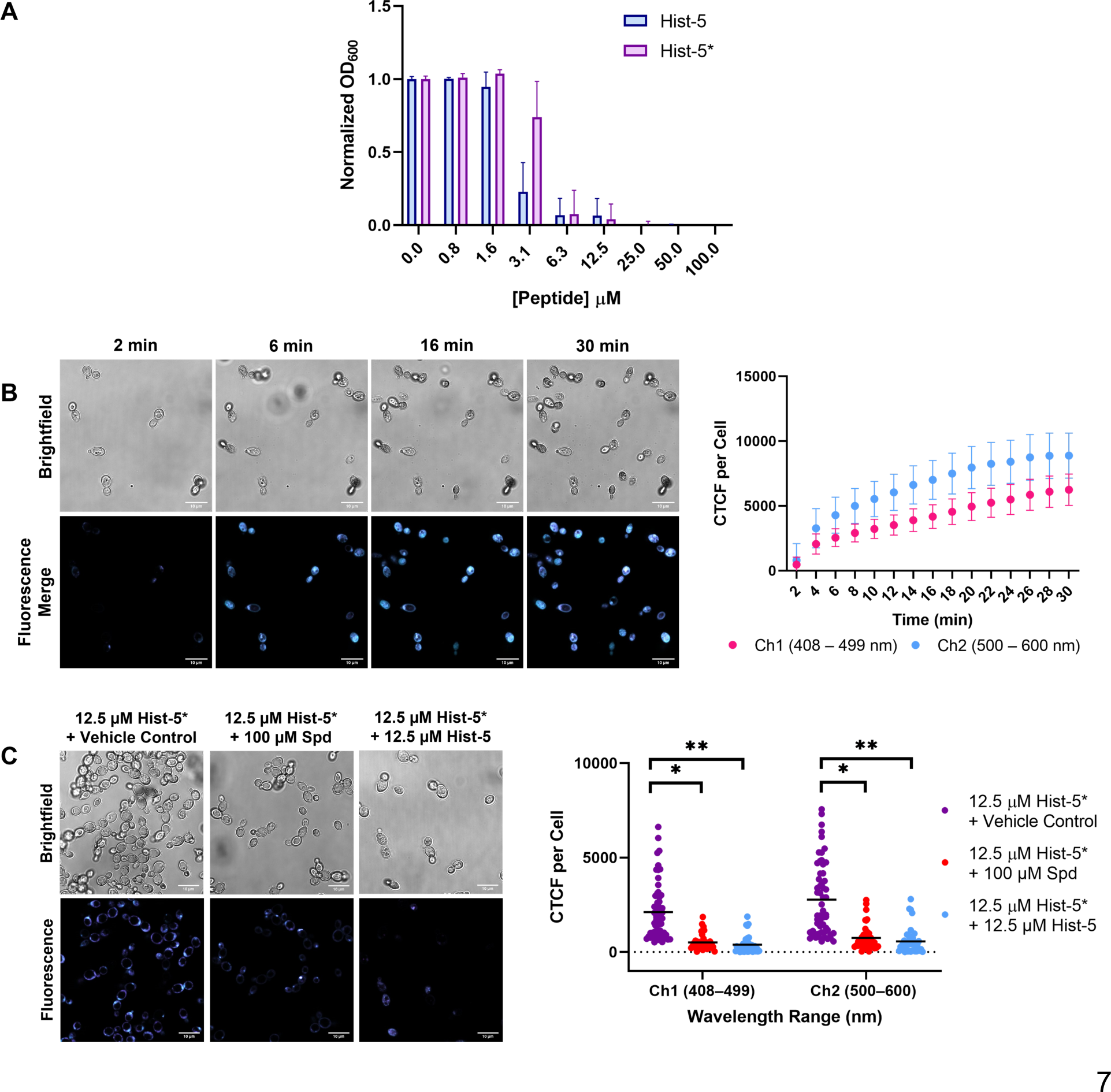
Hist-5* retains antifungal activity and uptake similar to native Hist-5. **(A)** *C. albicans* cells were pre-incubated with increasing concentrations of Hist-5 (blue) or Hist-5* (purple) for 1.5 h at 37 °C in PPB pH 7.4. Aliquots were resuspended in YPD and cell growth was measured by OD_600_ after 48 h incubation at 30 °C. **(B)** Timelapse microscopy images of cells treated with 12.5 µM Hist-5* at room temperature (RT) over 30 min in PPB. Corrected total cell fluorescence (CTCF) per cell is reported over time for each fluorophore channel. **(C)** Confocal fluorescence microscopy images of cells treated with Hist-5* + vehicle control (purple), + spermidine (red), or + Hist-5 (blue) as competitive substrates for cellular uptake. Cells imaged at RT for 5 min in PPB. CTCF for individual fluorescence channels under each treatment condition were quantified, each dot representing fluorescence values from individual cells on experiments carried out on three separate days. Error bars represent the standard deviation between three biological replicates. Scale bar = 10 µm. (* indicates p < 0.05, ** indicates p < 0.01, n = 3)

Confocal fluorescence microscopy was used to observe uptake of Hist-5* into fungal cells suspended in PPB at pH 7.4. Samples of *C. albicans*, in the yeast form, were treated with 12.5 µM Hist-5* and imaged over a thirty-minute period at room temperature (Figure 2B). Fluorescence emission from the Mca and ABD fluorophores was collected in separate wavelength channels, 1 and 2 respectively. Fluorescence from the two channels was found to colocalize, thus in all subsequent experiments the merged image of the two fluorescence channels is presented (Figure S5). Internalization of the labeled peptide into the cytosol was observed within five minutes of treatment (Figure 2B), thus in subsequent experiments cells were imaged over a five-minute time frame. Untreated cells and cells treated with 50 µM unlabeled Hist-5 displayed no detectable fluorescence indicating the observed fluorescence signal from cells treated with Hist-5* was not due to autofluorescence from the buffer, cells, or unlabeled peptide (Figure S6).

Hist-5 has been reported to use polyamine transporters Dur3/31 for intracellular translocation by *C. albicans* and cells grown in the presence of spermidine (Spd), the native substrate of these transporters, exhibit reduced uptake and killing activity of Hist-5.^10^ We therefore performed competition assays with Hist-5* and Spd to confirm that the labeled peptide still competes with Spd uptake by Dur3/31 (Figure 2C). Internalization of Hist-5* was measured by quantifying the corrected total cell fluorescence (CTCF) per cell from fluorescence microscopy images. We observed a statistically significant (p = 0.0116) decrease in Hist-5* uptake in cells treated with a combination of labeled peptide and Spd, compared to cells treated with Hist-5* and vehicle control. Unlabeled Hist-5 was also used as a competitive inhibitor for Hist-5* uptake and resulted in a significant (p = 0.0093) decrease in internalization of the labeled peptide (Figure 2C). These data demonstrate that our modified Hist-5* peptide utilizes the same polyamine transport system that native Hist-5 uses for uptake into fungal cells. Altogether, these data validate Hist-5* as a novel fluorescent analogue that retains antifungal activity and uptake mechanisms similar to native Hist-5.

### Zn^2+^ inhibits the antifungal activity of Hist-5 in a concentration- dependent manner

Two-dimensional broth microdilution checkerboard assays were performed to gain a more comprehensive understanding of the effect of Zn^2+^ on Hist-5 activity across a range of peptide and Zn^2+^ concentrations (Figure 3). In these experiments, cells suspended in PPB were first exposed to varying concentrations of Zn^2+^ and Hist-5 prior to incubation in a Zn-free synthetic defined medium (SD-Zn). Experiments were conducted in SD-Zn to rigorously control the exposure of cells to extracellular Zn^2+^, ensuring that their only exposure would be upon combination treatment of Hist-5 and Zn^2+^ in PPB. Under these rigorously-controlled conditions, Hist-5 alone exhibited potent antifungal activity against *C. albicans* cells, with an MIC of 0.8 µM in SD-Zn, indicating that Zn^2+^ is not required for Hist-5 activity (Figure 3). We note that this 0.8 µM MIC of Hist-5 against *C. albicans* cells in SD-Zn medium is significantly lower than that of the 25 µM MIC obtained in YPD medium, (Figures 2A and 3). This difference is likely because cells grown in a minimal media like SD-Zn are more sensitive than cells grown in a nutrient-rich broth like YPD. Interestingly, as cells were exposed to increasing concentrations of Zn^2+^, the MIC of Hist-5 increased. The most striking inhibitory effect of Zn^2+^ on Hist-5 occurred in cells exposed to the highest supplemental Zn^2+^ concentration, where the MIC of Hist-5 increased roughly 16-fold, from 0.8 µM to 12.5 µM (Figure 3). A similar Zn-induced inhibitory effect on antifungal activity was observed with our labeled Hist-5* peptide (Figure S7).

**Figure 3.**
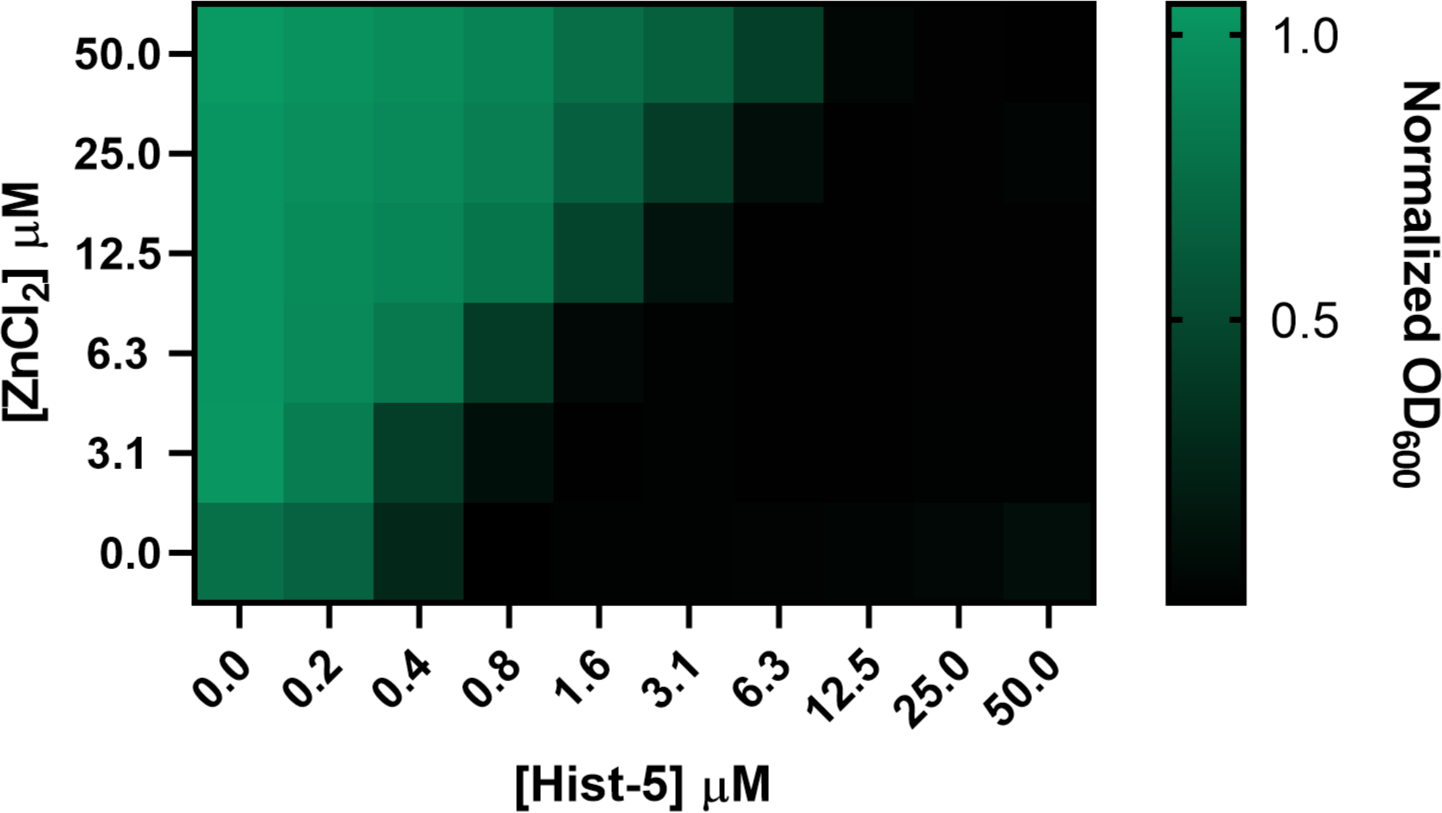
Fungicidal activity of Hist-5 is inhibited by increasing concentration of Zn^2+^. *C. albicans* cells were pre-incubated in PPB pH 7.4 for 1.5 h at 37 °C with increasing concentrations of Hist-5 and ZnCl_2_ as indicated in the figure axes. Aliquots were resuspended in 50 mM Tris-buffered synthetic defined Zn-free media (SD-Zn), pH 7.4, and cell growth was measured by OD_600_ after incubation for 48 h at 30 °C. Values represent the average from three separate biological replicates.

The Zn-induced reduction of Hist-5 activity observed in our experiments seemingly contradict previous studies that found Zn^2+^ increases Hist-5 activity.^25^ In order to reconcile these differences, we conducted checkerboard assays in which cells were exposed to low concentrations of Zn^2+^ and Hist-5 in a 1:2 ratio, mimicking the conditions described by Norris *et al.* in which they observed a Zn-induced increase in Hist-5 fungicidal activity.^25^ Indeed, when cells were treated with sub-inhibitory concentrations of Hist-5, Zn^2+^ supplementation resulted in an increase in peptide activity, with the strongest effects observed at ratios of 1:2 Zn^2+^ to peptide (Figure S8).

However, as the concentration of Zn^2+^ surpassed the concentration of Hist-5, the effects of Zn^2+^ switched from promoting Hist-5 fungicidal activity to inhibiting it (Figure S8). These results highlight the dynamic nature by which Zn^2+^ can modulate the antifungal activity of Hist-5, and help to reconcile seemingly contradictory conclusions.

Previous reports of Zn-induced peptide dimerization and aggregation were conducted using concentrations of Zn^2+^ and Hist-5 above 300 μM, significantly higher than the maximum peptide concentration of 50 μM used in our experiments.^24–27^ To determine whether Hist-5 dimers or aggregates were forming in our system, we used circular dichroism (CD) spectroscopy to monitor Hist-5 secondary structure in the presence of excess Zn^2+^ in both aqueous PPB and trifluoroethanol (TFE). Hist-5 exists in a random coil conformation in aqueous solution and adopts a more ordered, alpha-helical structure in membrane-like environments.^2, 43^ We observed a decrease in ellipticity upon addition of excess Zn^2+^ in both aqueous and organic solvent (Figure 4A). For Hist-5 in a random coil conformation, the ellipticity at 198 nm was plotted against increasing concentrations of Hist-5 in the presence of excess Zn^2+^ and fit to a linear model, R^2^ = 0.999 (Figure 4B), with ellipticity expected to vary linearly with concentration in the absence of oligomerization.^44^ For Hist-5 in an alpha-helical conformation, ellipticity at 222 nm was plotted against peptide concentration in the presence of excess Zn^2+^ and fit to a linear model, R^2^ = 0.998 (Figure 4C). We did not observe any deviations from linearity up to 50 µM Hist-5 in either PPB or TFE, indicating that under our conditions Zn-induced peptide dimerization/aggregation is not occurring.

**Figure 4.**
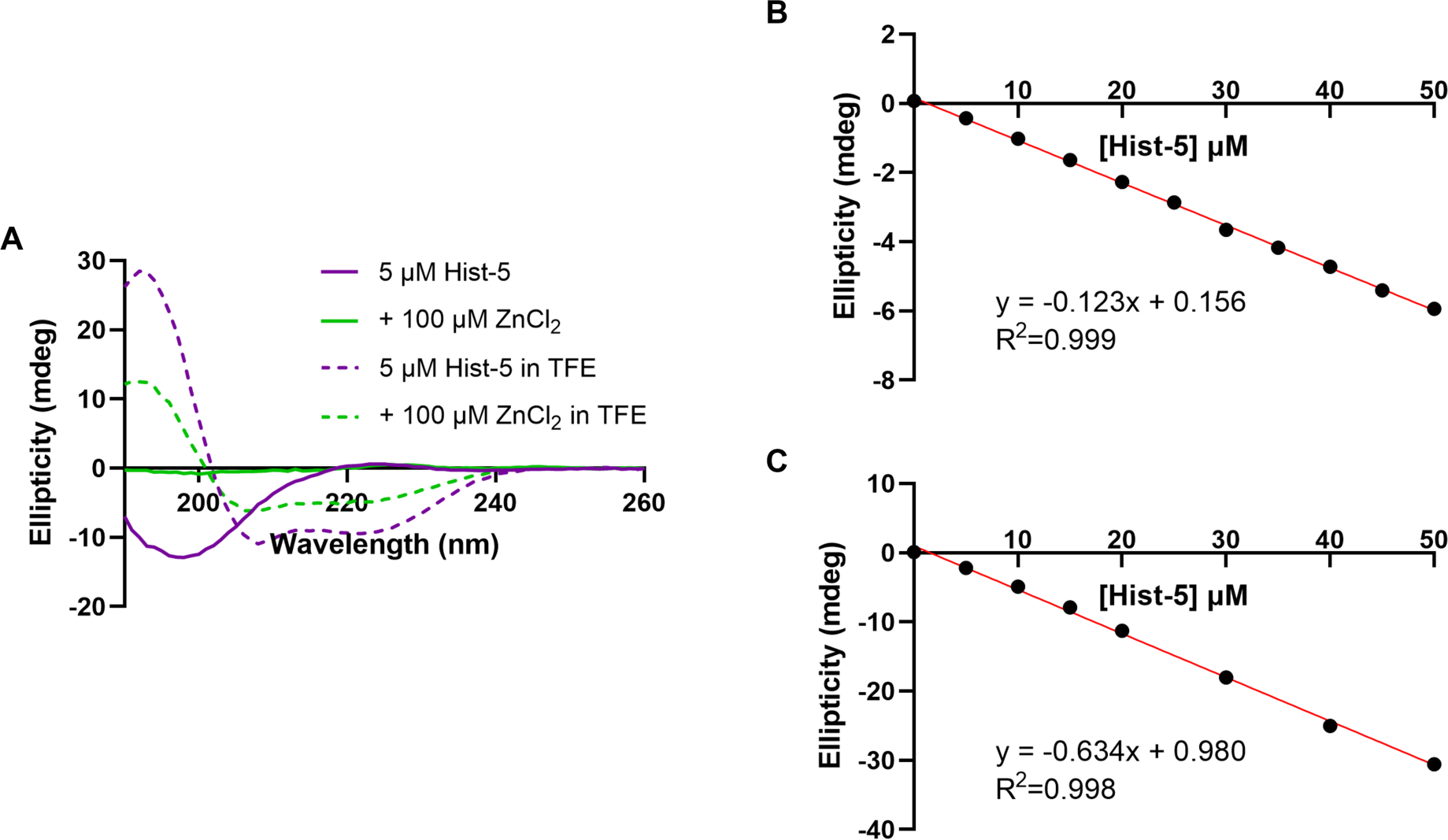
Changes in Hist-5 ellipticity remain linear up to 50 µM added peptide. **(A)** Full CD spectrum of 5 µM Hist-5 (purple) with or without 100 µM ZnCl_2_ (green) in PPB (solid lines) or 98% trifluoroethanol, TFE (dashed lines). **(B)** Titration of 0 – 50 µM Hist-5 into a solution containing 250 µM ZnCl_2_ in PPB. The ellipticity at 198 nm is plotted against peptide concentration and fit to linear model (R^2^ = 0.999). **(C)** Titration of 0 – 50 µM Hist-5 into a solution containing 250 µM ZnCl_2_ in 98% TFE. The ellipticity at 222 nm is plotted against peptide concentration and fit to linear model (R^2^ = 0.998). Data represent the average of three scans.

### Supplemental Zn^2+^ promotes surface adhesion of peptide to *C. albicans* and inhibits internalization

Uptake of Hist-5 by *C. albicans* is widely accepted as a requirement for antifungal activity, as Hist-5 is thought to have intracellular targets.^9, 25, 29, 41^ We observed a decrease in Hist-5 activity as a function of increasing Zn^2+^ concentration, thus we performed timelapse microscopy with *C. albicans* exposed to Hist-5* treated with a range of added Zn^2+^ concentrations to determine whether the decrease in antifungal activity could be the result of decreased peptide internalization. Rapid uptake and cytosolic fluorescence of Hist-5* in cells treated with peptide alone or peptide and a sodium chloride control were observed (Figure 5A). As the molar ratio of Zn^2+^ to peptide was increased from 0:1 to 0.5:1, there was reduced uptake of Hist-5* (Figure S9). The most striking results occurred in cells treated with 1:1 or 2:1 molar equivalents of Zn^2+^ to peptide. Under these conditions Hist-5* no longer appeared to internalize into the fungal cell, but instead remained bound to the cell surface (Figure 5A). This surface-bound state is evidenced in the localization of Hist-5* fluorescence around the perimeter of the cells, as shown in the intensity profiles (Figure 5A). The change in peak shape and the shift to a bimodal distribution directly correspond to the shift in peptide fluorescence and localization from the cytosol to the cell surface, as a function of Zn^2+^ concentration (Figure 5A). Adherence of Hist-5* to the cell perimeter was observed for up to two hours with timelapse microscopy (Figure S10), indicating that this effect is not transient. Additionally, this effect is specific for Zn^2+^ and does not occur in cells treated with equivalent concentrations of Hist-5* and other divalent metal cations, Cu^2+^ and Co^2+^ (Figure S11). The decrease in peptide internalization as a function of Zn^2+^ concentration was quantified and reported as CTCF (Figure 5B). A slight, but significant (p = 0.0245), decrease in Hist-5* internalization was seen in cells treated with 0.25 equivalents of Zn^2+^, while cells treated with 0.5 equivalents of Zn^2+^ or more experienced even greater decreases in peptide internalization (p < 0.001).

**Figure 5.**
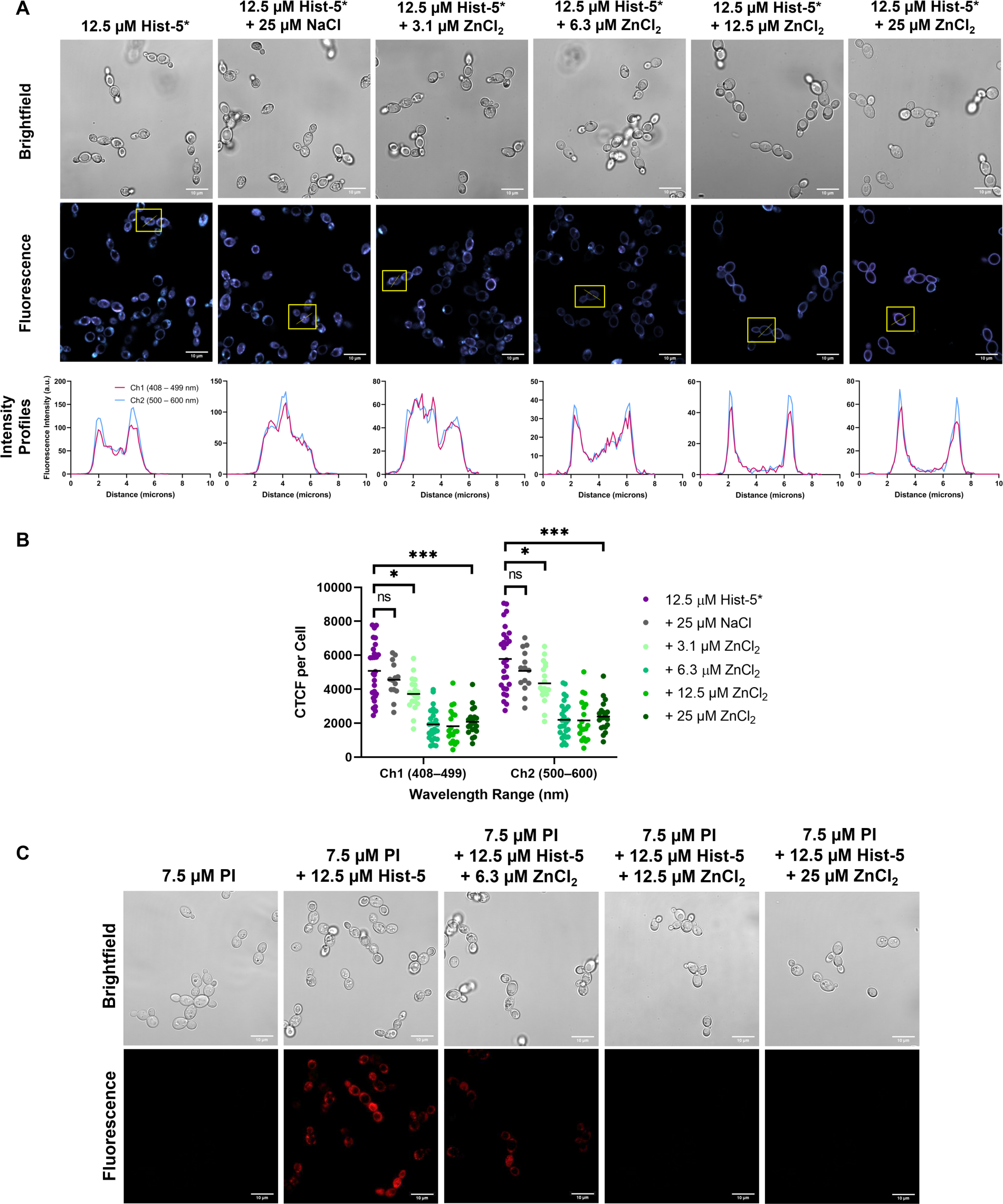
Zn^2+^ supplementation induces adhesion of peptide to the cell surface and inhibits peptide internalization and membrane activity. (A) Confocal fluorescence microscopy images of *C. albicans* cells treated with 12.5 µM Hist-5* alone, peptide+2 eq. NaCl, or varying concentrations of ZnCl_2_ (0.25, 0.5, 1, or 2 eq. Zn^2+^) at RT for 5 min in PPB. Fluorescence intensity profiles of representative cells (yellow box) under each treatment condition are plotted against distance in microns. (B) CTCF for individual fluorescence channels under each treatment condition were quantified; error bars represent the standard deviation between three separate biological replicates. (C) Confocal microscopy images of cells treated with 12.5 µM Hist-5, 7.5 µM PI, and varying concentrations of ZnCl_2_ (0.5, 1, or 2 eq. Zn^2+^) at RT for 5 min in PPB. Scale bar = 10 µm. (* indicates p < 0.05, *** indicates p < 0.001, n = 3)

To determine how Zn^2+^ supplementation affects Hist-5 membrane disruptive activity, we performed confocal fluorescence microscopy experiments with native Hist-5 peptide, using propidium iodide (PI) as a fluorescent indicator of membrane integrity (Figure 5C). Hist-5 induces membrane disruptions in *C. albicans* that allows leakage of PI into the cytosol. Treatment of cells with Hist-5 in combination with submolar equivalents of Zn^2+^ also resulted in internalization of PI, indicating membrane permeability. However, exposure of fungal cells to Hist-5 treated with one or more molar equivalents of Zn^2+^ did not cause dye leakage (Figure 5C), suggesting that these concentrations of Zn^2+^ protect against Hist-5-induced membrane disruption. Combined, our results show that Zn^2+^ supplementation alters Hist-5 uptake and activity by promoting peptide localization to the cell surface, leading to a decrease in peptide internalization, and inhibiting peptide membrane activity.

*C. albicans* is a polymorphic organism that transitions between yeast, pseudohyphal, and hyphal forms and the yeast to hyphae transition is critical for fungal virulence and pathogenesis ^45^. Formation of hyphae allows *C. albicans* to invade host epithelial and endothelial cells, causing infections, such as, oral thrush and vaginal candidiasis.^45^ Given the importance of *C. albicans* hyphae in pathogenesis, we also wanted to investigate the interactions between Hist-5 and the hyphal form of *C. albicans*. We observed internalization of Hist-5* in *C. albicans* hyphae treated with peptide alone and found that the addition of equimolar Zn^2+^ inhibited Hist-5* uptake, causing the peptide to adhere to the cell surface (Figure 6A). We also investigated the membrane activity of Hist-5 on *C. albicans* hyphae and found the Hist-5 alone was able to permeabilize the cell membrane and allow leakage of PI, however, Zn^2+^ supplementation provided a protective effect against Hist-5-induced membrane disruption (Figure 6B). These results are consistent with our observations of Zn-induced inhibition of peptide uptake and membrane activity in the yeast form of *C. albicans*.

**Figure 6.**
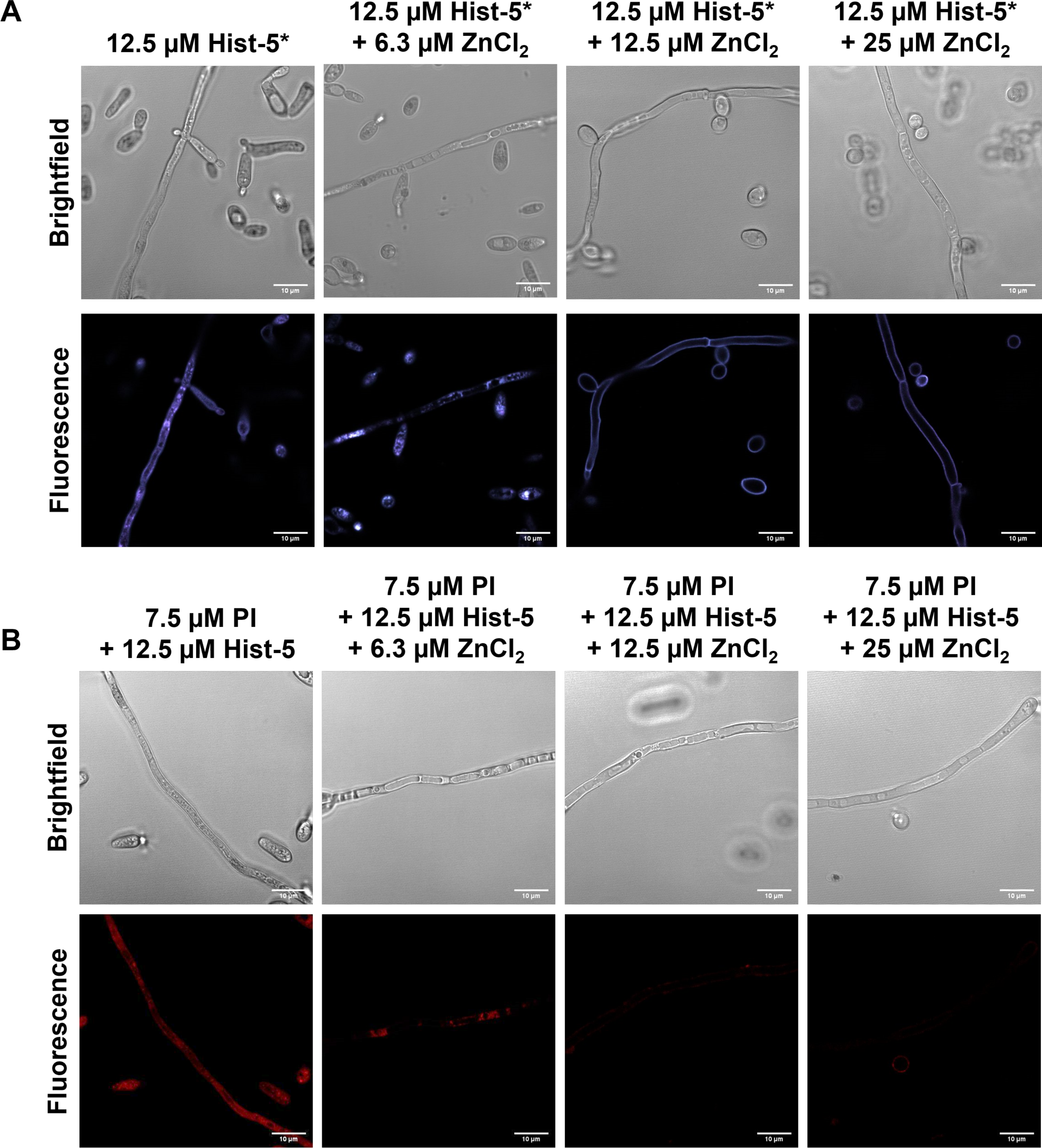
Zn^2+^ supplementation induces adhesion of Hist-5* to the cell surface and inhibits peptide internalization and membrane activity in the hyphal form of *C. albicans*. (A) Confocal fluorescence microscopy images of *C. albicans* cells in the hyphal form treated with 12.5 µM Hist-5* alone and varying concentrations of ZnCl_2_ (0.5, 1, or 2 eq. Zn^2+^) at RT for 5 min in PPB. **(B)** Confocal microscopy images of hyphal cells treated with 12.5 µM Hist-5, 7.5 µM PI, and varying concentrations of ZnCl_2_ (0.5, 1, or 2 eq. Zn^2+^) at RT for 5 min in PPB. Scale bar = 10 µm.

### A direct interaction between Zn^2+^ and Hist-5 results in peptide adhesion to the cell surface

Although we observed metal-dependent changes to Hist-5* fluorescence response in vitro (Figure 1D), these changes in the ABD:Mca ratio were not robust enough to detect with imaging. Therefore, we were unable to use this method to distinguish whether the surface-bound signal arose from direct peptide–Zn interaction at the cell surface. To determine whether Zn-induced surface adhesion of peptide to the fungal cell stems from a direct binding interaction between Hist- 5 and Zn^2+^, we used a Zn-responsive fluorophore to further interrogate the nature of the Zn–peptide interaction. Zinquin (ZQ) is a fluorescent Zn^2+^ sensor that can be used to detect labile cellular Zn^2+^ ions as well as protein-bound Zn^2+^.^46–48^ When ZQ detects labile Zn^2+^, a fluorescent Zn(ZQ)_2_ complex is formed which has an emission maximum centered around 500 nm. When ZQ forms an adduct with Zn-containing proteins, the emission spectrum undergoes a characteristic blue-shift to 480 nm.^47, 48^ Titration of Hist-5 into a solution of Zn(ZQ)_2_ in PPB resulted in a clear blue shift from 500 nm to 480 nm (Figure S12), indicating that Hist-5 binds to Zn^2+^ in a manner that enables simultaneous binding of Zn^2+^ and ZQ to form a ternary complex in vitro.

Whole cell fluorescence spectroscopy experiments with *C. albicans* cells were performed to determine whether ZQ-Zn-Hist-5 complexes also form in a more complex cellular environment. Figure 7A shows fluorescence emission spectra of a suspension of cells in PPB exposed to various combinations of ZQ, Zn^2+^ and Hist-5. Addition of ZQ alone to cells shows minimal background fluorescence which increases significantly upon addition of Zn^2+^, with an emission at 500 nm indicating formation of the Zn(ZQ)_2_ complex, as expected. While the addition of Hist-5 alone did not affect the background ZQ emission, cells treated with a combination of Hist-5, ZQ, and increasing amounts of Zn^2+^ resulted in an increase in the fluorescent signal. This increase in fluorescence emission was accompanied with a blue shift in the emission spectra, with cells exposed to a 1:1 or greater Zn-to-peptide molar ratio exhibiting the strongest blue shift (Figure 7A), indicating an interaction between ZQ, Zn and Hist-5.

**Figure 7.**
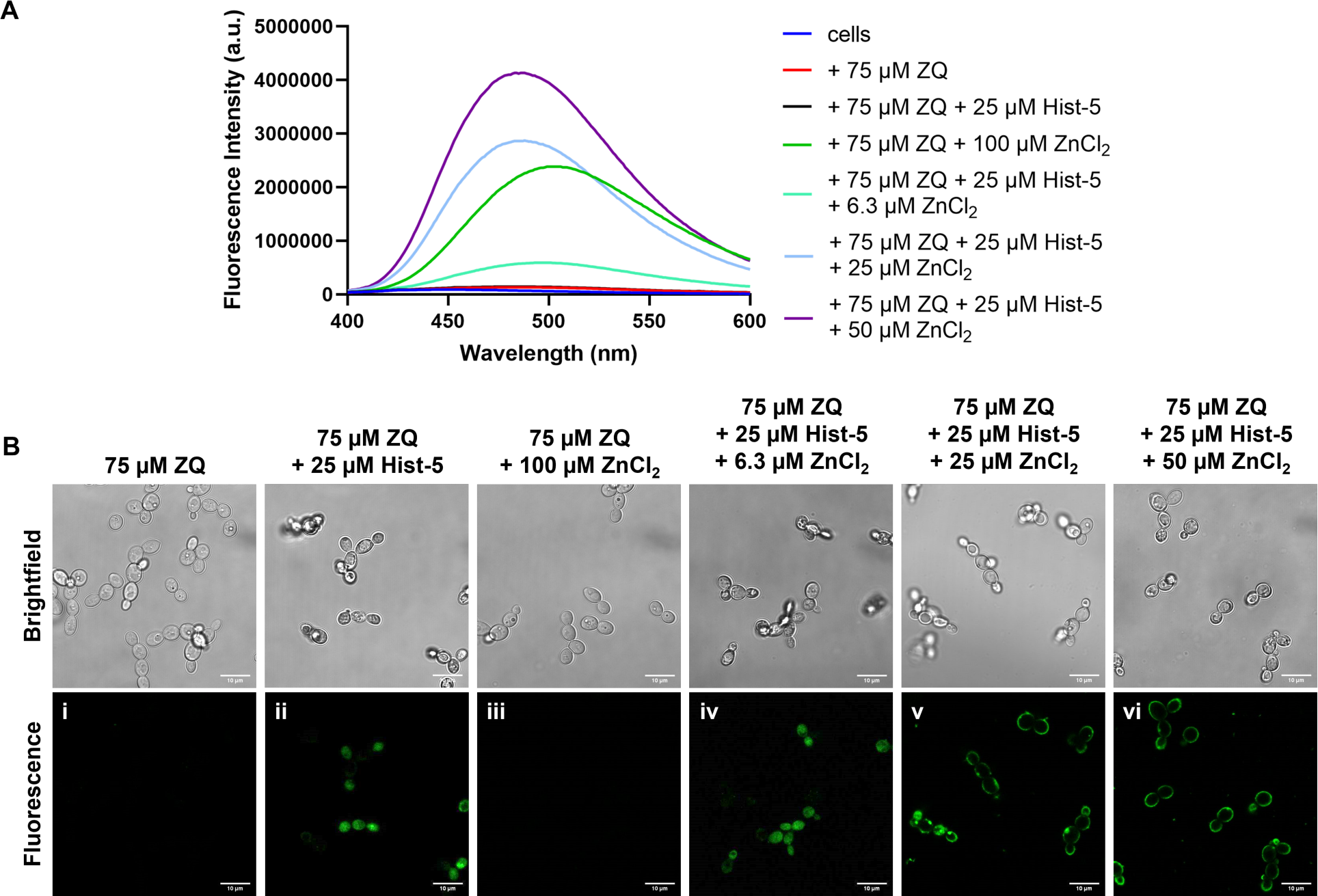
Zn^2+^ binding to Hist-5 promotes adhesion to the cell surface. **(A)** Titration of ZnCl_2_ into a solution containing 75 µM zinquin acid (ZQ), 25 µM Hist-5, and 200 µL *C. albicans* (∼10^6 cells) in PPB pH 7.4. **(B)** Confocal fluorescence microscopy images of cells treated with ZQ+ Zn^2+^, ZQ+Hist-5, and ZQ+Hist-5 and a variety of ZnCl_2_ concentrations (0.25, 1, and 2 eq. Zn^2+^) at RT for 5 min in PPB. Scale bar = 10 µm.

Complementary confocal microscopy studies were also performed, using the same concentrations and treatment conditions established in our whole cell spectrofluorometry assays to investigate the location of ZQ-responsive Zn^2+^ as a function of Hist-5 treatment in *C. albicans*.

Although cell suspensions treated with ZQ and Zn^2+^ exhibited a strong fluorescence signal in vitro, there was no detectable fluorescence in the microscopy images of cells treated with ZQ either alone or in combination with Zn^2+^ (Figure 7B, panels i and iii). The acid form of ZQ used in these experiments is known to have poor membrane permeability, so these results indicated that the probe, even in the presence of added Zn^2+^, does not internalize or otherwise interact with cells in a way that would produce a detectable and localized Zn-responsive signal. However, fluorescence emission was observed in the cytosol of cells treated with Hist-5 and ZQ, as well as those treated with Hist-5, ZQ, and submolar equivalencies of Zn^2+^ relative to peptide (Figure 7B, panels ii and iv). This observation could result from the membrane-disruptive effects of Hist-5, which could enable ZQ permeability and subsequent detection of intracellular accessible Zn^2+^. In cells treated with ZQ and equimolar or higher ratios of Zn^2+^ to peptide, however, fluorescence was distinctly localized around the cell perimeter (Figure 7B, panels v and vi). This change in localization of the fluorescence response under equimolar Zn^2+^ conditions is reminiscent of the changes observed in *C. albicans* cells treated with the labeled Hist-5* peptide and Zn^2+^. While the microscopy images alone cannot distinguish between Zn(ZQ)_2_ complexes and Zn-ZQ-P ternary complexes with proteins or peptides, the combination of the fluorescence and imaging data with ZQ combined with the results from labeled Hist-5* provide strong evidence of a direct binding interaction between Hist-5 and Zn^2+^ that changes the recognition and uptake of Hist-5 into *C. albicans* by restraining the peptide to the cell surface.

### Modulation of extracellular Zn^2+^ concentration by metal-binding molecules reverses Zn-induced surface adhesion of Hist-5

Taken together, our data show that the effects of Zn^2+^ on Hist-5 activity, uptake, and localization are dependent on Zn^2+^ concentration in the surrounding environment. In order to test whether Zn-induced binding of Hist-5 to the cell surface could be reversed by decreasing Zn^2+^ availability, cells were initially exposed to peptide treated with one molar equivalent of Zn^2+^ to induce adhesion to the cell surface, then subsequently exposed to a Zn^2+^ chelating molecule. Cells were monitored over time for peptide internalization (Figure 8). Addition of the extracellular metal chelator ethylenediaminetetraacetic acid (EDTA) led to recovery of Hist-5* internalization, as evidenced by the change in peptide localization from the cell perimeters to the cytosol (Figure 8A, left). In addition, Zn^2+^ chelation away from Hist-5 by EDTA resulted in the peptide regaining its membrane disruptive activity, allowing PI leakage into the fungal cell (Figure 8A, right). EDTA has been known to cause membrane permeability on its own,^49^ however under our conditions, PI leakage can be attributed solely to Hist-5 membrane activity, as PI uptake into the cytosol was not observed in cells treated with these concentrations of EDTA and PI (Figure S13).

**Figure 8.**
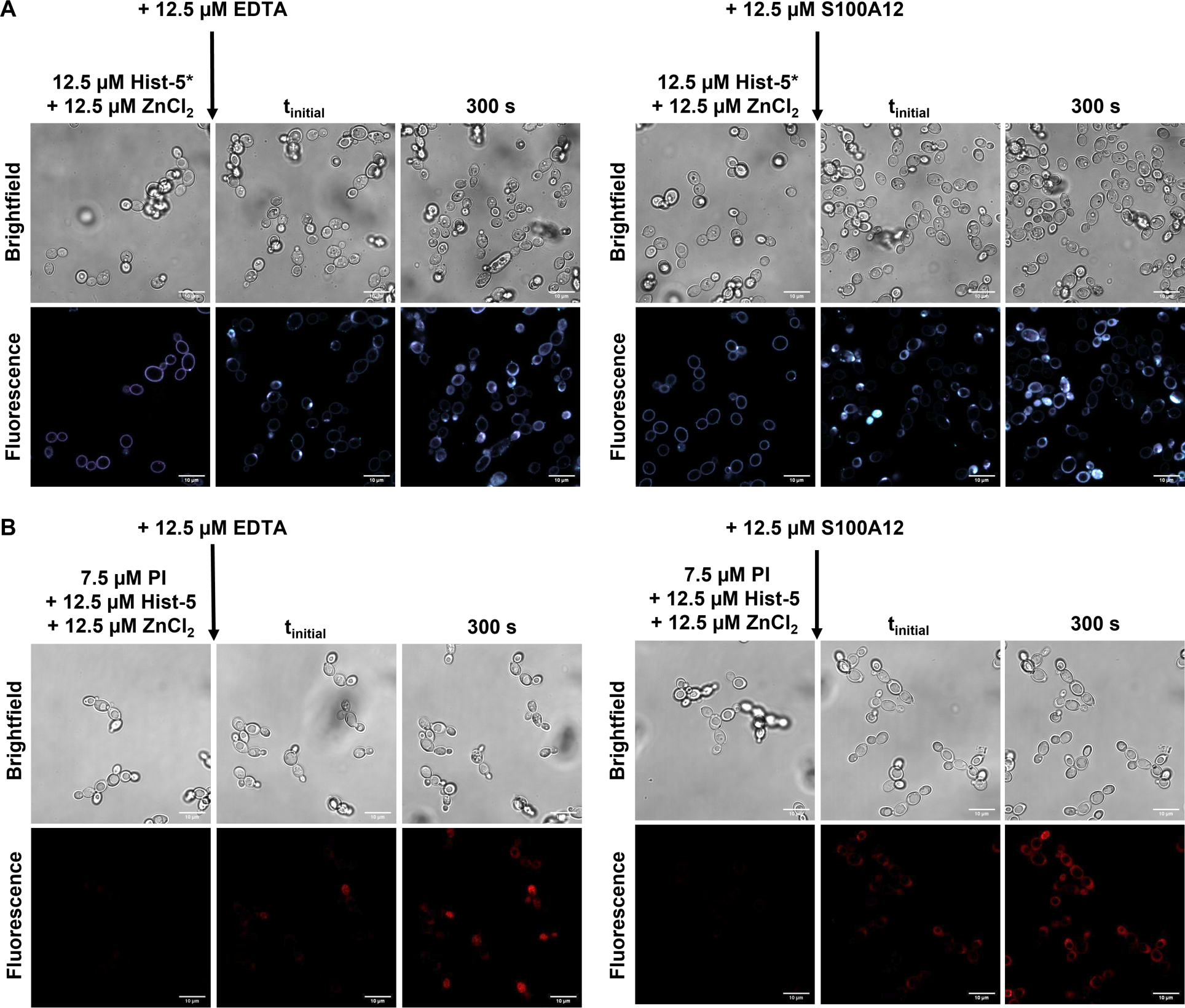
Addition of Zn^2+^ binding chelators or proteins reverses adhesion of peptide to the cell surface. **(A)** Timelapse microscopy images of *C. albicans* cells treated with 12.5 µM Hist-5*+1 eq. ZnCl_2_ or 12.5 µM Hist-5+7.5 µM PI and 1 eq. ZnCl_2_ for 2.5 min at RT in PPB, followed by addition of 12.5 µM EDTA, as indicated with arrow, images were collected over 300 s at RT in PPB. **(B)** Timelapse microscopy images of cells treated with 12.5 µM Hist-5*+1 eq. ZnCl_2_ or 12.5 µM Hist-5+7.5 µM PI and 1 eq. ZnCl_2_ for 2.5 min at RT in PPB, followed by addition of 12.5 µM S100A12, as indicated with arrow, images were collected over 300 s at RT in PPB. Scale bar = 10 µm.

In order to verify whether the results with EDTA could be replicated with more biologically relevant conditions, treated cells were also exposed to S100A12, a human host-defense protein that binds Zn^2+^ with high affinity^50, 51^ and is released during infection.^52^ Addition of S100A12 to cells treated with Hist-5* and an equivalent of Zn^2+^ led to a reversal of Zn-induced surface binding of the peptide, resulting in uptake and internalization of Hist-5* (Figure 8B, left). Treatment of fungal cells with S100A12 also led to the recovery of membrane permeabilization activity by Hist- 5 and PI uptake (Figure 8B, right). These results parallel the changes in peptide internalization and activity that were observed with EDTA treatment and have interesting implications for how Hist- 5 may operate in the context of the wider immune response.

## Discussion

Although the role of Zn^2+^ on Hist-5 antifungal activity has previously been investigated, probing how Hist-5 operates across a range of Zn^2+^ concentrations allowed us to gain a full account of its effect on peptide activity. Throughout our studies we utilized Hist-5* to visualize peptide internalization and localization within the fungal cell. We did not observe granular intracellular distribution of Hist-5*, in contrast to a prior report of fluorescein-labeled Hist-5 which attributed the staining effect to Hist-5 localization to mitochondria.^41^ Instead, we observed uniform cytosolic distribution of Hist-5* along with an apparent buildup of Hist-5* at a localized point along the cell surface, reminiscent of the spatially restricted sites observed in a separate report of a fluorescein-labeled Hist-5 by Mochon *et al.*^42^ Through our experiments, we found that Zn^2+^ availability greatly affected Hist-5 antifungal activity and internalization. Our data demonstrate that submolar ratios of Zn^2+^ to peptide improve Hist-5 antifungal activity and allow for peptide internalization (Figures 3, 5, and S8). However, as the concentration of Zn^2+^ increases, Hist-5 antifungal activity is inhibited and its uptake is blocked (Figures 3 and 5). It is likely this concentration-dependent effect of Zn^2+^ has resulted in the varied reports regarding the effect of Zn^2+^ on Hist-5 activity in the literature.^20, 25, 28, 29^ While our data reveal clear Zn-dependent effects on Hist-5 activity and uptake, the question that remains is why does Zn^2+^ affect Hist-5 in this manner?

*C. albicans* is a commensal organism that inhabits mucosal membranes of the human body, including the oral cavity. Under immunocompetent host conditions, *C. albicans* living in the oral environment are constantly exposed to varying levels of Hist-5,^31, 35^ raising a question about the function of this immunopeptide beyond antifungal cell killing. In a healthy individual, is Hist-5 actively entering and killing commensal microbial cells, or is there a surveillance mechanism that triggers the antifungal response when the microbial environment is disrupted and infection is initiated? Here, we add to the growing body of literature suggesting that Hist-5 may participate in interactions with cells that are not solely for antifungal purposes but rather promote and maintain microbial homeostasis for oral microbial health.^29, 53^

We propose a working model for the role of Zn^2+^ in Hist-5 antifungal activity and commensalism where modulation of exchangeable Zn^2+^ concentration in the growth environment acts as a dial to tune Hist-5 uptake and activity in *C. albicans* (Figure 9). When the host is healthy, commensal *C. albican*s are continuously exposed to sublethal concentrations of Hist-5 and Zn^2+^. This constant exposure causes the cells to become less virulent and exhibit a stress-adapted response in which they promote an anti-inflammatory signaling from oral epithelial cells by altering the composition of cell wall polysaccharides.^29^ We hypothesize that at these concentrations of Zn^2+^ and Hist-5, Zn-bound peptide adheres to the cell surface, but does not internalize and exert antifungal activity against the commensal *C. albicans* cells. However, the switch from Hist-5 coexisting with commensal *C. albicans* to Hist-5 employing its antifungal activity could be triggered by a change in environmental Zn^2+^ concentrations, perhaps among other triggers. We showed that the inhibitory effects of Zn^2+^ on Hist-5 uptake and membrane activity were reversible by adding an extracellular chelating agent to decrease the amount of Zn^2+^ available to the peptide (Figure 8). In this model we suggest that when the host is fighting an infection that signals *C. albicans* to switch from commensal to pathogenic, a change must also be registered by Hist-5 to start exerting its antifungal properties. We posit that Hist-5 switches to act as an antifungal agent when there is a decrease in salivary Zn^2+^ availability, as the result of the body engaging the processes of nutritional immunity.^38^ While high affinity metal chelating proteins like S100A12 are being deployed at the host-pathogen interface to deplete surrounding Zn^2+^ levels,^52^ they also chelate Zn^2+^ away from Hist-5 enabling the peptide to exert its antifungal activity (Figure 9). Hist-5 may then work to kill the pathogenic fungal cell in two ways: either by permeabilizing the cell membrane (Figure 8)^25^ or by internalizing (Figure 8) and exhibiting the classical Hist-5 antifungal mode of action.^39^

**Figure 9.**
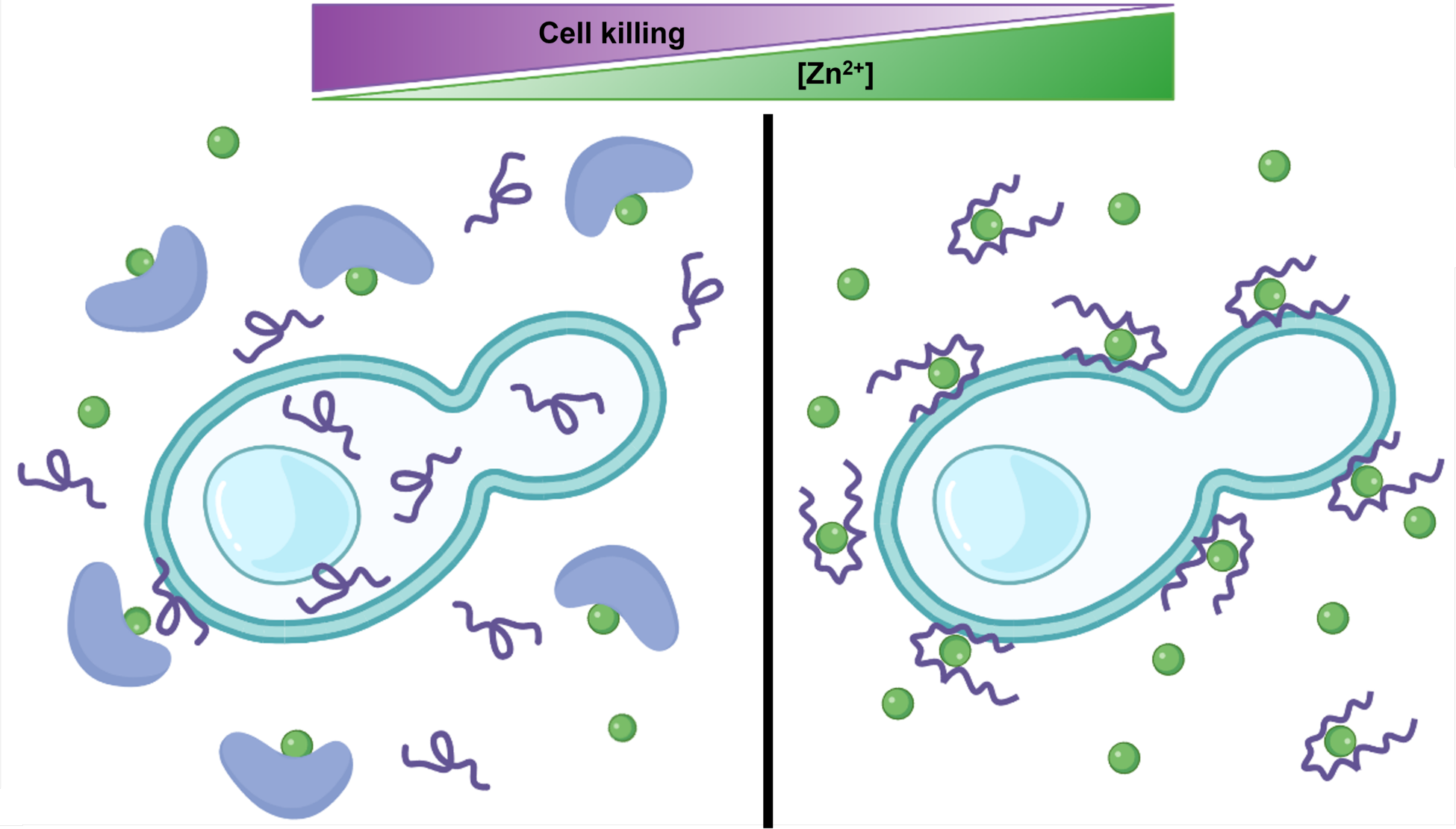
Modulation of exchangeable Zn^2+^ in the growth environment acts as a dial to tune Hist-5 antifungal activity. Graphical representation of proposed Hist-5- Zn^2+^ interactions and how they affect peptide internalization and activity. When Hist-5 is exposed to *C. albicans* cells in a Zn-replete environment, the peptide binds Zn^2+^ (green spheres), adheres to the cell surface, and does not exhibit antifungal activity against the cell. When the concentration of Zn^2+^ in the growth environment decreases due to chelation of Zn^2+^ by host defense proteins (blue protein), Zn^2+^ is removed from Hist-5, and the peptide internalizes into the cell allowing Hist-5 to exert its antifungal activity. Figure created with BioRender.com.

In conclusion, here we demonstrate a novel role for Zn^2+^ as a regulatory switch that can either be used to maintain commensalism between Hist-5 and *Candida* or induce antifungal activity. Our data offer context for how Hist-5 and its interactions with metals operate within the larger immune response at the host-pathogen interface.

## Experimental section

### Materials and general methods

Chemicals and solvents were obtained from commercial suppliers and used as received unless otherwise noted. All aqueous solutions were prepared using Milli-Q water. Stock solutions were prepared either in DMSO or Milli-Q water.

Stock solutions of Cu^2+^ (45 mM), Fe^3+^ (100 mM), Mn^2+^ (100 mM), Zn^2+^ (10 mM), and Co^2+^ (10 mM) were prepared from CuSO_4_·5H_2_O, FeCl_3_·6H_2_O, MnCl_2_·4H_2_O, ZnCl_2_, and CoCl_2_ in Milli-Q water. Stock solutions of propidium iodide (PI, Sigma-Aldrich) and zinquin free acid (ZQ, 2- methyl-8-(toluene-p-sulfonamido)-6-quinolyloxyacetic acid, Santa Cruz Biotechnology Inc.) were prepared at 5 mg/mL in DMSO, aliquoted, frozen, and stored in the dark when not in use. Working solutions of spermidine (Spd, Acros Organics), 50 mg/mL in Milli-Q water, were prepared fresh on the day of the experiment. S100A12 homodimer protein was generously provided by Prof. Elizabeth Nolan’s Lab of Massachusetts Institute of Technology.

### Peptide synthesis

Peptides were synthesized on a Protein Technologies PS3 automated peptide synthesizer using rink amide MBHA low-loaded resin (Protein Technologies) on a 0.1 mmol scale. All Fmoc- protected amino acids were purchased from Chem-Impex International Inc. unless otherwise stated. Amino acid coupling was achieved using O-benzotriazole-N,N,N′,N′- tetramethyluronium hexafluorophosphate (HBTU, Protein Technologies) in the presence of N-methylmorpholine as an activator in N,N′-dimethylformamide (DMF) for 30 min cycles. 20% piperidine in DMF was used to deprotect Fmoc groups during the synthesis. Prior to cleavage, the resin was washed three times with 1 – 2 mL each of glacial acetic acid, then dichloromethane, followed by methanol. Side chain deprotection and peptide cleavage from the resin were achieved by treatment with 5 mL of a solution of 95% trifluoroacetic acid (TFA), 2.5% ethanedithiol, and 2.5% triisopropylsilane (TIS, Sigma Aldrich) for 3.5 h under N_2_ gas to yield peptides with N-terminal free amines and amidated C-termini. A continuous flow of N_2_ gas was used to evaporate TFA to a volume of 2 mL. Afterward, the peptide was precipitated and washed three times with diethyl ether, and dried in air prior to purification.

### Synthesis of Hist-5*

Doubly-labeled Hist-5* was prepared in two steps, starting with singly labeled Hist-5Mca, which was synthesized by incorporating Fmoc-beta-(7- methoxycoumarin-4-yl)-Ala-OH (Mca, Bachem) at position 10 and Fmoc-Cys(Trt)-OH (Novabiochem) at position 24 via solid phase peptide synthesis, as described above. Purified and quantified Hist-5Mca was then reacted under basic conditions in a 1:1 molar ratio with 27.3 µM 4-fluoro-7-sulfamoylbenzofurazan (ABD-F, TCI America) in DMSO for 20 min over a 60 °C water bath. Peptides were purified using a Waters 1525 reverse-phase Binary High-Performance Liquid Chromatography (HPLC) Pump on a Waters XBridge Prep C18 Column (10 µm OBD, 19 mm × 250 mm) with a 40 m linear gradient from 3 to 97% acetonitrile to water, with 0.1% TFA. Purity was validated to >95% using HPLC on a Waters XBridge Peptide BEH C18 Column (130Å, 10 µm, 4.6 mm × 250 mm) and the masses of the peptides were confirmed by electrospray ionization mass spectrometry (ESI-MS).

Hist-5 sequence DSHAKRHHGYKRKFHEKHHSHRGY, calculated mass: 3034.5, found (M + 6H^+^) 506.9 (Figure S2).

Hist-5Mca sequence: DSHAKRHHG(Mca)KRKFHEKHHSHRGC, calculated mass: 3056.5, found (M + 4H^+^) 765.1 m/z (Figure S3).

Hist-5* sequence DSHAKRHHG(Mca)KRKFHEKHHSHRG(ABD) calculated mass: 3253.5, found (M + 4H^+^) 814.4 m/z (Figure S4).

### Quantification of peptide stock solutions

Peptide stock solutions were prepared by dissolving ∼0.05 g lyophilized peptide in 1 mL of Milli-Q water. The concentration of stock solutions was determined using the Edelhoch method.^54^ In short, 4−6 μL of peptide stock was diluted into 400 μL of 8 mM urea to obtain an absorbance at 278 nm between 0.1 and 1 absorbance unit. Absorption spectra were recorded in 1 cm quartz cuvettes on a Varian Cary 50 UV−vis spectrophotometer. The concentration of Hist-5 stock solution was determined from the A_278_ readings using an extinction coefficient of 1450 M^−1^ cm^−1^ for each tyrosine.^55^ The concentration of Hist-5Mca and Hist-5* stock solutions were determined from the A_325_ readings using an extinction coefficient of 12,000 M^-1^ cm^-1^ for methoxycoumarin.^56^ Peptide stock solutions were stored at -20 °C in sealed cryogenic storage vials.

### Fluorescence spectroscopy

#### Metal-dependent changes to Hist-5* fluorescence

Fluorescence spectra were collected in a 5 mm quartz Starna Micro fluorometer cell using an Edinburgh Instruments FS5 Fluorometer.

Emission spectra for Hist-5* were collected over 412 – 600 nm with an excitation wavelength of 405 nm, using 2.0/2.0 nm excitation/emission bandwidths. Fluorescence of Hist-5* in PPB was monitored as a function of increasing equivalents of ZnCl_2_ or NaCl as an anion control. The fluorescence emission from the two fluorophores were identified by wavelength ranges from 412 – 499 nm for Mca and 500 – 600 nm for ABD. The sum of the fluorescence signal for each fluorophore as well as the ratio between the ABD and Mca fluorophores signals were plotted as a function of equivalents of metal added to visualize the metal-dependent changes to Hist-5* fluorescence

#### In vitro formation of ZQ-Zn(II) complexes

Fluorescence spectra were collected in 5 mm quartz Starna Micro fluorometer cell using an Edinburgh Instruments FS5 Fluorometer. Emission spectra were collected over 400 – 600 nm with an excitation wavelength of 370 nm, using 2.0/2.0 nm excitation/emission bandwidths. Increasing equivalents of Hist-5 were titrated into a solution containing 25 µM ZQ and 12.5 µM ZnCl_2_ in PPB, with a final volume of 200 µL in the cuvette. For whole cell fluorescence experiments, *C. albicans* were prepared in the same manner used for microscopy experiments, described above. Increasing equivalents of ZnCl_2_ were added into a solution containing 75 µM ZQ, 25 µM Hist-5, and 200 µL *C. albicans* (∼10^6 cells) in PPB.

#### Circular dichroism (CD) spectroscopy

All CD spectra were collected using an AVIV Model 435 CD spectrometer with a 1 nm bandwidth at 25 °C. The full CD spectra of 5 µM Hist-5 and 5 µM Hist-5+100 µM ZnCl_2_ in PPB or 98% trifluoroethanol (TFE) were collected in a 1 cm quartz cuvette from 260 – 190 nm. Scans were taken using 1 nm steps with a 6 s averaging time. Data reported are the average of three scans. For titration experiments of 0 – 50 µM Hist-5 in the presence of 250 µM ZnCl_2_ in PPB or 98% TFE, scans were collected in kinetics mode using a 1 mm or 1 cm quartz cuvette for Hist-5 in PPB or TFE, respectively. Scans were taken at 198 nm for Hist-5 in PPB and 222 nm for Hist-5 in TFE, with a 1 s averaging time over 60 s. The ellipticity at 198 nm was plotted against peptide concentration and fit to a linear model in GraphPad Prism (R^2^ = 0.999). Data reported are the average of 60 scans.

#### Synthetic defined (SD) media

All tris-buffered synthetic defined media formulations were prepared from Chelex-treated Milli-Q water with individual addition of media components to allow for rigorous control of metal content. To deplete trace metals from water prior to media preparation, Milli-Q water was treated with Chelex 100 resin 100–200 mesh sodium form via batch method (50 g/L, Bio-Rad Laboratories). A concentrated stock of SD media not containing Cu^2+^, Fe^3+^, Mn^2+^, or Zn^2+^ (10× SD-) was prepared in the Chelex-treated MilliQ water by adding glucose and yeast nitrogen base (YNB) ingredients at 10× concentrations. YNB components were added individually to avoid trace metals present in commercial YNB mixtures. To prepare working 1X SD medium, 10× SD- was diluted into Chelex-treated water (1:10), and Ultra-Pure Tris–HCl (VWR) was added to a final concentration of 50 mM. The pH of the media was adjusted to 7.4 using 1.0 M HCl or NaOH pellets, this media was then filter-sterilized. Finally, CuSO_4_, FeCl_3_, MnCl_2_, and ZnCl_2_ were added, as appropriate to create either individual metal dropout or metal complete media (SD+).

#### Yeast strains and culture conditions

Fungal stocks were maintained in 25% glycerol in YPD at −80 °C. Experiments were performed with *C. albicans* clinical isolate SC5314, which was obtained from the American Type Culture Collection (ATCC). Prior to all experiments, *C. albicans* were streaked onto yeast peptone dextrose (YPD, Gibco) agar plates from frozen glycerol stocks and incubated at 30 °C for 24 h. A single colony was used to inoculate 5 mL YPD or SD+ media, which was then incubated at 30 °C, 200 rpm overnight for 16 – 18 h or 24 h, respectively, to stationary growth phase.

### Cellular growth inhibition assays

#### Microdilution assays

*C. albicans* were cultured overnight in YPD, as described above, and diluted to an optical density at 600 nm absorption (OD_600_) of 0.07 in PPB pH 7.4 and used as the working culture. Peptides to be tested were serially diluted 2-fold in PPB from aqueous stocks and plated in a clear, flat-bottomed 96-well plate. 100 μL of the working culture of cells were then added to the 96-well plate, containing PPB and peptide, to a final OD_600_ of 0.035 and a final volume of 200 µL per well. Final concentrations of peptide in the plate are indicated in the figure axes. For each experiment, a peptide-free positive growth control and a cell-free, negative control were included. This plate was incubated for 1.5 h at 37 °C, 200 rpm to allow time for peptide to interact with cells. After incubation, 10 μL aliquots from the plate were added to a new 96-well plate containing 190 μL YPD media with a final volume of 200 μL per well. The new media plate was then incubated for 48 h at 30 °C, 200 rpm. All media plates were covered with air-permeable AeraSeal film (Sigma) to minimize evaporation. Fungal growth was evaluated via OD_600_ using a PerkinElmer Victor3 V multilabel plate reader at 0, 24, and 48 h. OD_600_ values were normalized to the positive growth control and adjusted by subtracting the 0 h timepoint readings from other timepoint data at 24 and 48 h, to remove any background signal from YPD. Data are representative of three biological replicates, each with three technical replicates per experiment. For a single experiment, each of the three replicate conditions were averaged and the error was calculated as standard deviation, which is indicated by error bars in the figures. Final 48 h timepoint data is reported by plotting OD_600_ readings versus peptide concentration.

#### Two-dimensional broth microdilution checkerboard assays

*C. albicans* were cultured overnight in SD+ media, as described in the yeast strains and culture conditions section and diluted to an OD_600_ of 0.07 in PPB pH 7.4 and used as the working culture. Peptides to be tested were serially diluted from aqueous stocks 2-fold in PPB, right to left along the row in a clear, flat-bottomed 96-well plate. ZnCl_2_ was serially diluted from aqueous stock in water, down the column of the plate. Aliquots of 180 μL of the working culture of cells were then added to the 96-well plate, containing PPB, peptide, and Zn^2+^, to a final OD_600_ of 0.06 and a final volume of 200 µL per well. Final concentrations of peptide and Zn^2+^ in the plate are indicated in the figure axes. This plate was incubated for 1.5 h at 37 °C, 200 rpm. After incubation, 10 μL aliquots from the plate were added to three new 96-well plates containing 190 μL of SD-Zn or SD+ media with a final volume of 200 μL per well. The new SD-Zn or SD+ media plates were covered with AeraSeal film and incubated for 48 h at 30 °C and fungal growth measurements were taken as described above in the microdilution assays section. OD_600_ values were normalized to the positive growth control and adjusted by subtracting the 0 h timepoint readings from other timepoint data at 24 and 48 h, to remove any background signal from the media. Data are representative of three biological replicates, each with three technical replicates per experiment. To visualize the results, a final heatmap was generated in GraphPad Prism using average OD_600_ values from the biological replicates at 48 h. Concentrations of peptide and Zn^2+^ indicated in the figure represent the amount of peptide and Zn^2+^ present in the preincubation plate.

### Confocal fluorescence microscopy

#### Preparation of *C. albicans* in the yeast form for microscopy

*C. albicans* were cultured overnight in YPD, as described in the yeast strains and culture conditions section. Cells from the overnight culture were then diluted either 1:100 or 1:50 in 5 mL fresh YPD media. The subculture was allowed to grow to an OD_600_ of 1.0 at 30 °C, 200 rpm. Cells were pelleted at 5000 rpm for 20 min to remove excess media and then resuspended in PPB pH 7.4 for imaging.

#### Preparation of *C. albicans* in the hyphal from for microscopy

*C. albicans* were cultured overnight for 24 h in SD+ medium. After 24 h, cells in the overnight culture were diluted to an OD_600_ of 0.1 in fresh SD+ medium. The diluted cells were then aliquoted into two equal portions of 2 mL each. To induce hyphal formation, one of the subcultures was treated with 12.5 mM N-Acetylglucosamine (GlcNAc), the second subculture was left untreated as a control. The two subcultures were grown at 37 °C, 200 rpm for a further 24 h. Cells from the +GlcNAc and untreated control cultures were pelleted at 5000 rpm for 20 min to remove excess media. Cells from the two cultures were then resuspended in PPB pH 7.4 for imaging.

#### General microscopy parameters and setup

All experiments were performed using live cells, in the yeast form (unless otherwise stated) at room temperature, suspended in PPB, and plated into an ibidi µ-Slide 18 well to a final volume of 40 μL per well. Confocal images were acquired with a Zeiss 880 Ariyscan Inverted Confocal microscope using Plan Apochromat 63x/1.4 oil objectives. For most experiments, images were taken in 25 s intervals from 0 – 5 min, unless otherwise indicated. Images were obtained as 10–20 optical slices per wavelength spaced 0.5 µm apart along the Z-axis. Images presented are cells in the middle image of a Z-stack after 5 min of treatment with peptide, metal or dye, and are representative of cells in experiments conducted on three separate days. For experiments involving Hist-5*, the 405 nm diode laser was used to excite the two fluorophores. Fluorescence was detected over 408 – 499 nm for Mca using channel 1 and 500 – 600 nm for ABD using channel 2. Generally, Fiji software was used for image acquisition and processing. MATLAB ver.R2021b (MathWorks Natick, MA) software was used to conduct bulk image intensity analysis and determine corrected total cell fluorescence (CTCF) values per cell for all images, the full MATLAB script may be accessed in the supporting information. Each dot in the fluorescence intensity plots represents CTCF values of individual cells in images from experiments performed on separate days.

#### Hist-5* uptake and internalization over time

*C. albicans* were prepared for microscopy as described in the preparation of *C. albicans* for microscopy section. Cells were treated with 12.5 μM Hist-5* and uptake of the peptide was monitored via timelapse microscopy over 30 min with images being taken every 2 min. MATLAB software was used to perform image intensity analysis and plot CTCF intensity per cell for each channel over time. Error bars represent the standard deviation in CTCF values from experiments performed on three separate days.

#### Spermidine (Spd) Hist-5 competition assays

*C. albicans* were prepared for microscopy as described in the preparation of *C. albicans* for microscopy section. Cells were treated with 12.5 μM Hist-5* and either 100 μM Spd, 12.5 μM Hist-5 or an equal volume of PPB as a vehicle control.

Uptake of Hist-5* is shown in the microscopy images after 5 min and MATLAB software was used to perform image intensity analysis and plot CTCF intensity for each treatment condition. Statistical differences in CTCF values between treatments were calculated using an ordinary one-way ANOVA with Dunnett’s multiple comparison test in GraphPad Prism.

#### Effects of Zn^2+^ addition on Hist-5* and Hist-5 uptake and membrane activity

*C. albicans* were prepared for microscopy as described in the preparation of *C. albicans* for microscopy section. Cells were treated with 12.5 μM Hist-5* and the desired concentrations of ZnCl_2_ or NaCl as an anion control. Uptake of Hist-5* is shown in the microscopy images after 5 min and MATLAB software was used to perform image intensity analysis and plot CTCF intensity for each treatment condition. Statistical differences in CTCF values between treatments were calculated using an ordinary one-way ANOVA with Dunnett’s multiple comparison test in GraphPad Prism. Fiji software was used to generate intensity profiles for individual cells in an image. For membrane permeability assays, cells were treated with 7.5 μM PI, 12.5 μM Hist-5, and the desired concentrations of ZnCl_2_. Uptake of PI is shown after 5 min treatment. PI was excited using the 488 nm line of the argon ion laser and fluorescence was detected over 600 – 700 nm.

#### Zn-dependent changes in Hist-5* and Hist-5 uptake and membrane activity in the *C. albicans* hyphal form

Hyphal formation was induced in *C. albicans* and the cells were prepared for microscopy as described in the preparation of *C. albicans* in the hyphal from for microscopy section. Cells were treated with 12.5 μM Hist-5* and the desired concentrations of ZnCl_2_ and uptake of Hist-5* is shown in the microscopy images after 5 min. For membrane permeability assays, cells were treated with 7.5 μM PI, 12.5 μM Hist-5, and the desired concentrations of ZnCl_2_. Uptake of PI is shown after 5 min treatment. PI was excited using the 488 nm line of the argon ion laser and fluorescence was detected over 600 – 700 nm.

#### Fluorescence microscopy of ZQ-Zn(II) complexes

*C. albicans* were prepared for microscopy as described in the preparation of *C. albicans* for microscopy section. Cells were treated with 75 μM ZQ, 25 μM Hist-5 and the desired concentrations of ZnCl_2_. Fluorescence signals from ZQ complexes are shown after 5 min in the microscopy images. ZQ was excited with a 405 nm diode laser and fluorescence was detected over 420 – 600 nm.

#### Zinc chelation assays

*C. albicans* were prepared for microscopy as described in the preparation of *C. albicans* for microscopy section. Cells were initially treated with either 12.5 μM Hist-5* or Hist-5+PI and 12.5 μM ZnCl_2_. Peptide and Zn^2+^ treated cells were imaged for 2.5 min via timelapse microscopy to visualize surface adhesion of the peptide to the fungal cell. These cells were then exposed to 12.5 μM of a metal chelating agent, either EDTA or S100A12 protein and the cells were imaged via timelapse microscopy for a further 5 min. Uptake of either Hist-5* or PI after 5 min treatment with the chelating agent is shown in the microscopy images. PI was excited using the 488 nm line of the argon ion laser and fluorescence was detected over 600 – 700 nm.

## Supporting Information

The Supporting Information is available free of charge.

- Synthetic and peptide characterization details, Figures S1–S13 showing additional fluorescence microscopy images, spectra, growth assays, and MATLAB script.

## Author Contributions

JXC and KJF conceived and designed the experiments. JXC carried out the experiments. SG performed CD experiments, KSA wrote and implemented MATLAB script for image analysis. JXC and KJF analyzed the data and wrote the manuscript, with contributions from all coauthors.

## Supporting information

Campbell Supporting Information

## Acknowledgements

The antifungal and biological work presented here was supported by the National Institutes of Health (Grant R01GM084176), with funding from the National Science Foundation supporting the metal-binding chemistry (NSF CHE-1808710). We acknowledge the Duke Light Microscopy Core Facility, with funding from the shared instrumentation grant (1S10RR027867-01). We thank Prof. Elizabeth Nolan’s Lab at Massachusetts Institute of Technology for generously providing the S100A12 homodimer protein, and Dr. Catherine Denning-Jannace for help with hyphal cells.

## Abbreviations

Histatin-5, Hist-5; fluorescent histatin-5, Hist-5*; methoxycoumarin, Mca; sulfamoylbenzofurazan, ABD; potassium phosphate buffer, PPB; room temperature, RT; spermidine, Spd; optical density at 600 nm, OD_600_; minimum inhibitory concentration, MIC; corrected total cell fluorescence, CTCF; synthetic defined media, SD; circular dichroism, CD; trifluoroethanol, TFE; propidium iodide, PI; zinquin free acid form, ZQ; ethylenediaminetetraacetic acid, EDTA; solid phase peptide synthesis, SPPS; electrospray ionization mass spectrometry, ESI-MS.

