## Supplementary material for "Zinc binding inhibits cellular uptake and antifungal activity of Histatin-5 in *Candida albicans*": Campbell Supporting Information

**Figure S1. Synthesis of Hist-5*.**

**Figure S2. HPLC and MS Characterization of Hist-5.**

**Figure S3. HPLC and MS Characterization of Hist-5Mca.**

**Figure S4. HPLC and MS Characterization of Hist-5*.**

**Figure S5. Fluorescence signals from Mca and ABD fluorophores on Hist-5* colocalize.**

**Figure S6. Control cells exhibit no fluorescence signal.**

**Figure S7. Fungicidal activity of Hist-5* is inhibited by increasing concentration of $Zn^{2+}$.**

**Figure S8. Modulation of cell growth by $Zn^{2+}$ at sub-MIC levels of Hist-5, revealing that $Zn^{2+}$ improves Hist-5 activity at a ratio of 1:2 Zn:peptide under these conditions.**

**Figure S9. $Zn^{2+}$ addition blocks peptide internalization.**

**Figure S10. Zn-induced adherence of Hist-5* to the cell surface is long-lived.**

**Figure S11. Surface-bound effect of Hist-5* is Zn-specific.**

**Figure S12. In vitro formation of a ternary complex between Hist-5, ZQ, and $Zn^{2+}$.**

**Figure S13. EDTA and S100A12 protein do not induce membrane permeability.**

**MATLAB script for image intensity analysis.**

#### Figure S1. Synthesis of Hist-5*.

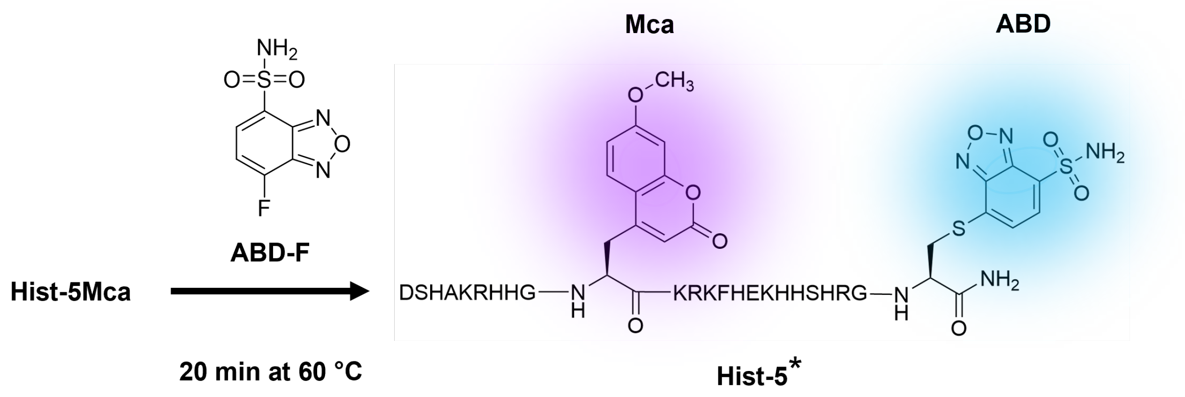


**Figure S1. Synthesis of Hist-5*.** Fmoc-protected methoxycoumarin (Mca) was incorporated into the solid phase peptide synthesis of Hist-5 yielding a singly labeled peptide, Hist-5Mca. Purified Hist-5Mca was reacted in a 1:1 molar ratio with 4-Fluoro-7-sulfamoylbenzofurazan (ABD-F) for 20 min in a water bath at 60 °C yielding the doubly labeled Hist-5 peptide, Hist-5*.

**Figure S2. HPLC and MS Characterization of Hist-5.**

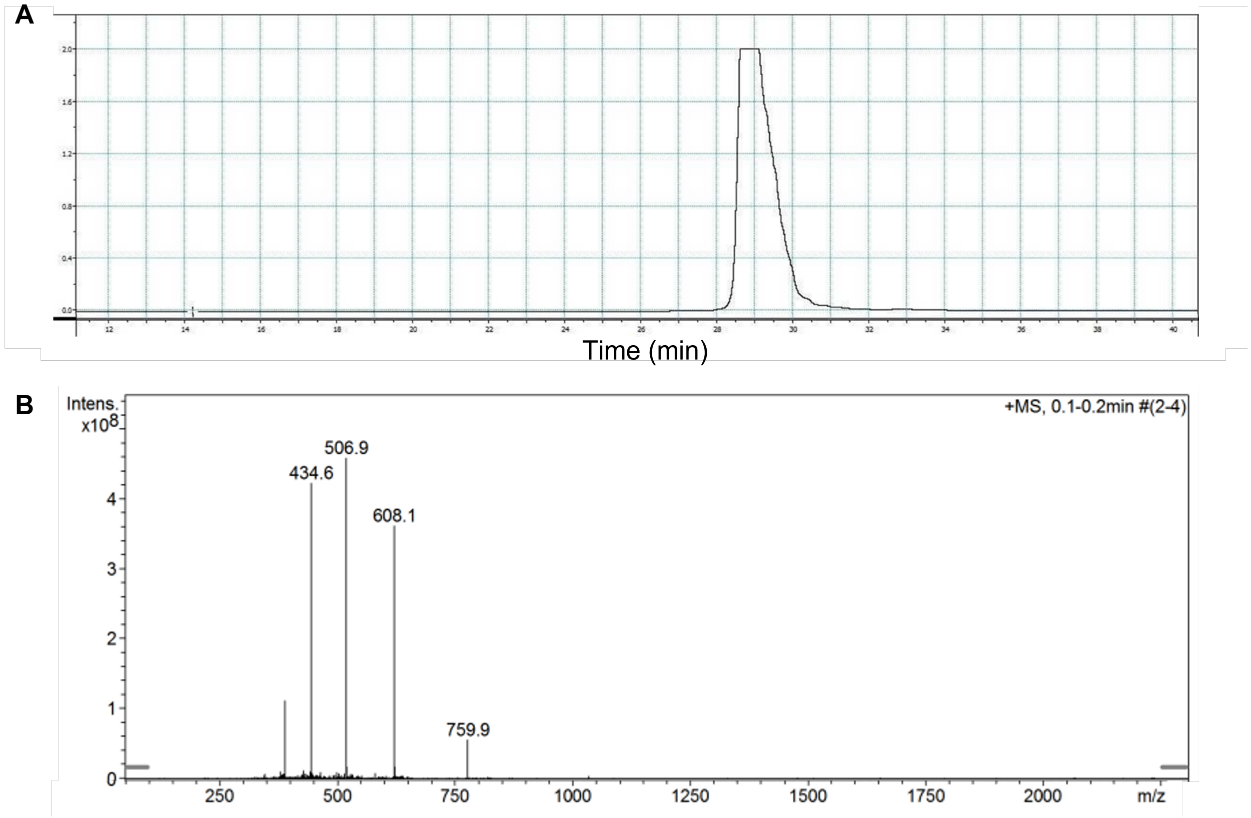


**Figure S2. HPLC and MS Characterization of Hist-5. (A)** Analytic HPLC trace of Hist-5 detection at 280 nm, retention time ~28.8 min. **(B)** Mass spectrum of Hist-5. Observed masses confirmed by electrospray ionization mass spectrometry: calculated mass for Hist-5 3034.5, found $(M + 4H^+)$ 759.9 m/z, $(M + 5H^+)$ 608.1 m/z, $(M + 6H^+)$ 506.9 m/z, $(M + 7H^+)$ 434.6 m/z.

### Figure S3. HPLC and MS Characterization of Hist-5Mca.

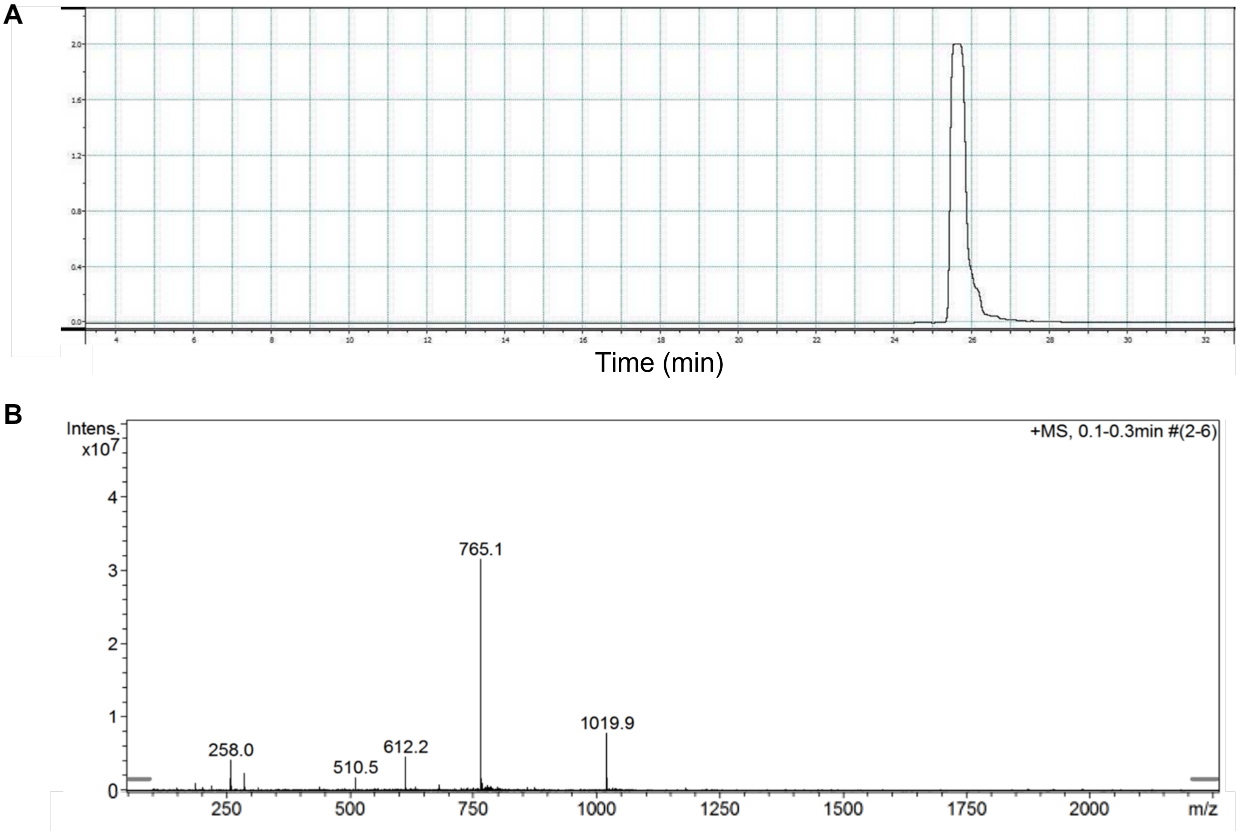


**Figure S3. HPLC and MS Characterization of Hist-5Mca. (A)** Analytic HPLC trace of Hist-5Mca detection at 325 nm, retention time ~25.7 min. **(B)** Mass spectrum of Hist-5Mca. Observed masses confirmed by electrospray ionization mass spectrometry: calculated mass for Hist-5-Mca 3056.5, found (M + $3H^+$) 1019.9 m/z, (M + $4H^+$) 765.1 m/z, (M + $5H^+$) 612.2 m/z, (M + $6H^+$) 510.5 m/z.

#### Figure S4. HPLC and MS Characterization of Hist-5*.

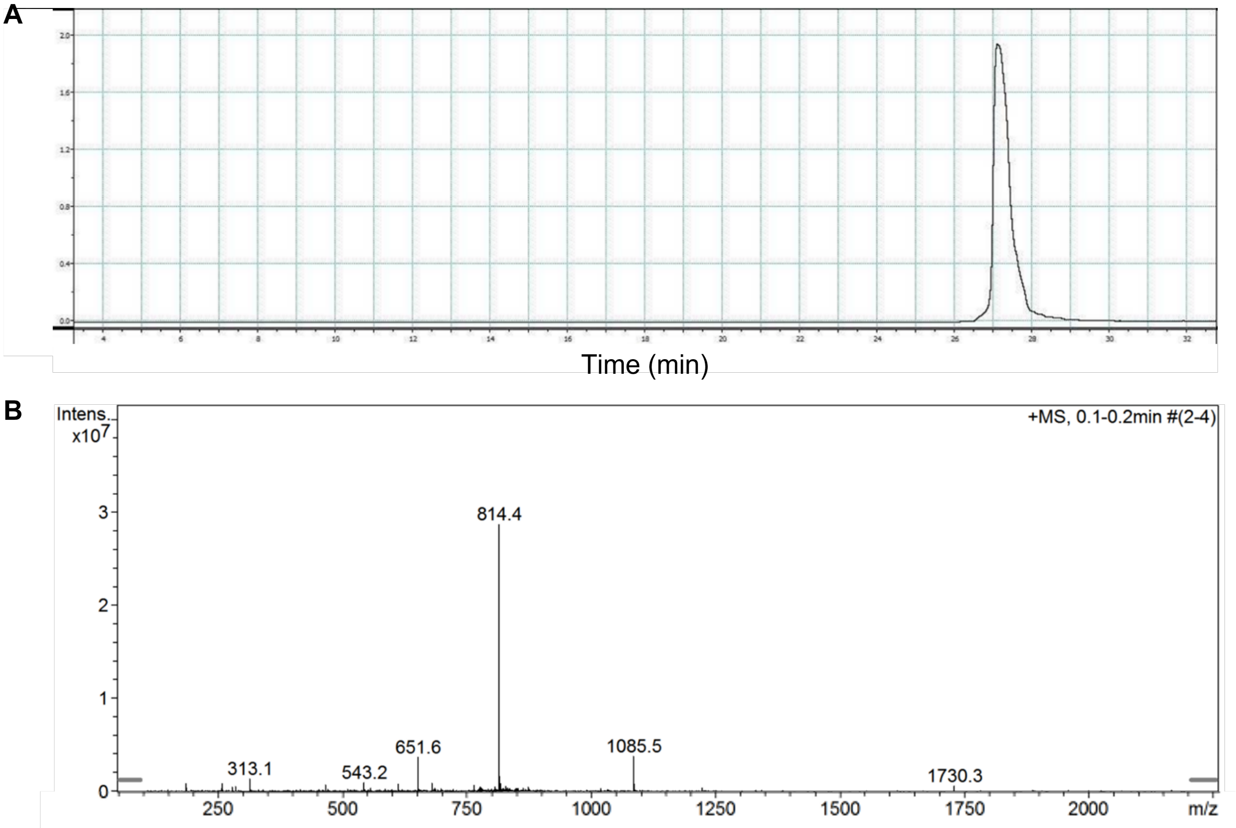


**Figure S4. HPLC and MS Characterization of Hist-5*. (A)** Analytic HPLC trace of Hist-5* detection at 325 nm, retention time ~27.1 min. **(B)** Mass spectrum of Hist-5*. Observed masses confirmed by electrospray ionization mass spectrometry: calculated mass for Hist-5* 3253.5, found (M + $3H^+$) 1085.5 m/z, (M + $4H^+$) 814.4 m/z, (M + $5H^+$) 651.6 m/z, (M + $6H^+$) 543.2 m/z.

**Figure S5. Fluorescence signals from Mca and ABD fluorophores on Hist-5* colocalize.**

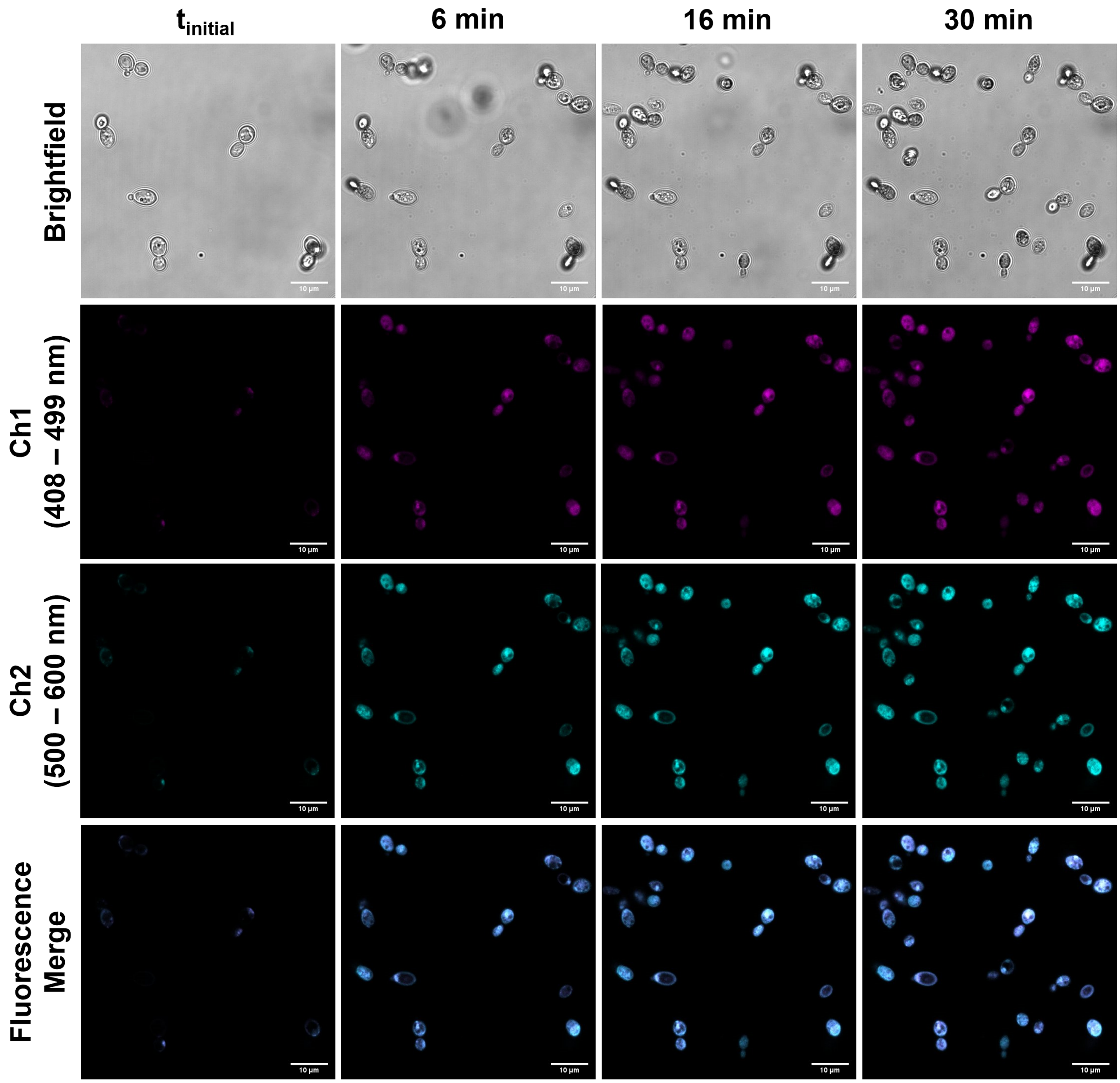


**Figure S5. Fluorescence signals from Mca and ABD fluorophores on Hist-5* colocalize.** Timelapse microscopy images of cells treated with 12.5 μM Hist-5* at room temperature (RT) over 30 min in 1 mM potassium phosphate buffer pH 7.4 (PPB). Scale bar = 10 μm.

**Figure S6. Control cells exhibit no fluorescence signal.**

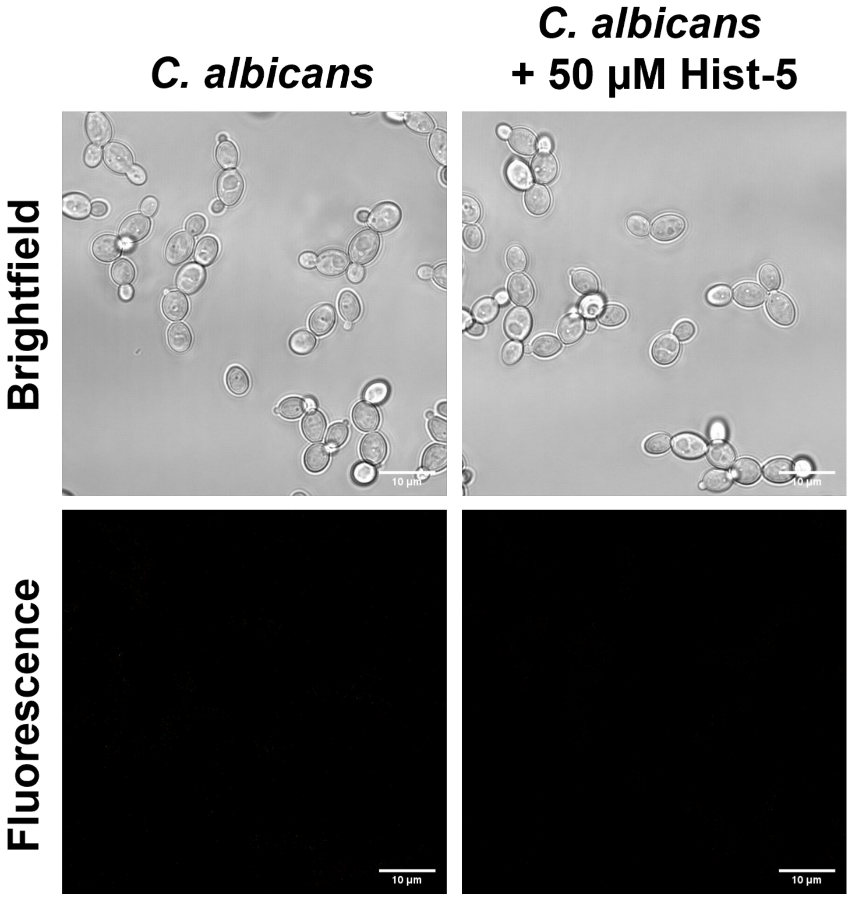


**Figure S6. Control cells exhibit no fluorescence signal.** Confocal fluorescence microscopy images of untreated *C. albicans* cells and cells treated with 50 µM Hist-5 at RT for 5 min in PPB.

**Figure S7. Fungicidal activity of Hist-5* is inhibited by increasing concentration of $Zn^{2+}$.**

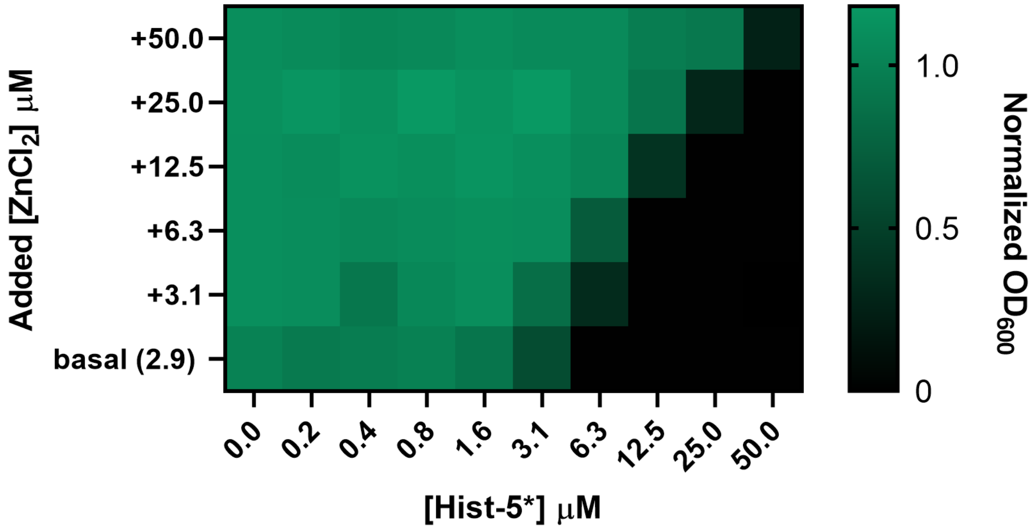


**Figure S7. Fungicidal activity of Hist-5* is inhibited by increasing concentration of $Zn^{2+}$.** *C. albicans* cells were pre-incubated in PPB pH 7.4 for 1.5 h at 37 °C with increasing concentrations of Hist-5* and supplemental $ZnCl_2$; concentrations are indicated in the figure axes. Aliquots were resuspended in 50 mM Tris-buffered synthetic defined media (SD+), pH 7.4, and cell growth was measured by $OD_{600}$ after incubation for 48 h at 30 °C. Values represent the average from three separate biological replicates.

**Figure S8. Modulation of cell growth by $Zn^{2+}$ at sub-MIC levels of Hist-5, revealing that $Zn^{2+}$ improves Hist-5 activity at a ratio of 1:2 Zn:peptide under these conditions.**

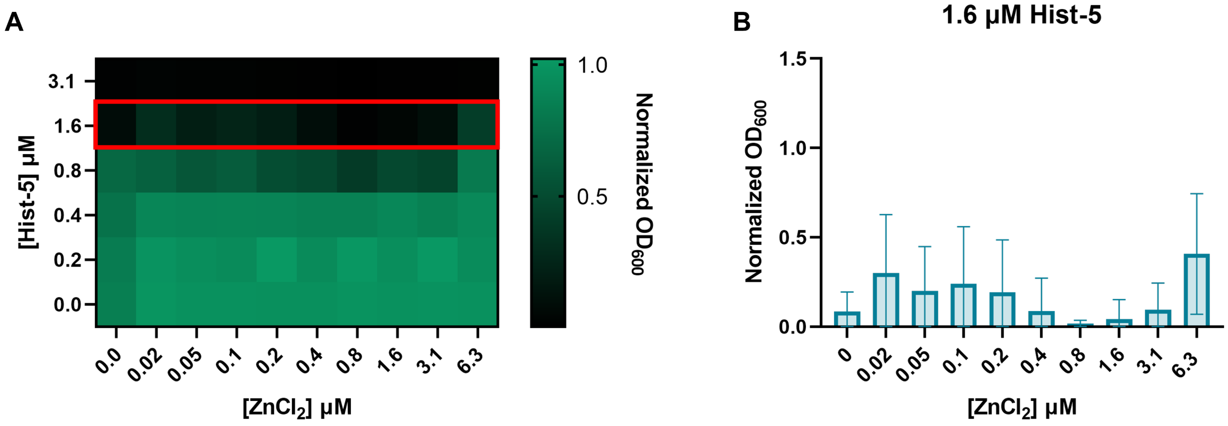


**Figure S8. Modulation of cell growth by $Zn^{2+}$ at sub-MIC levels of Hist-5, revealing that $Zn^{2+}$ improves Hist-5 activity at a ratio of 1:2 Zn:peptide under these conditions. (A)** *C. albicans* cells were pre-incubated in PPB pH 7.4 for 1.5 h at 37 °C with increasing concentrations of Hist-5 and supplemental $ZnCl_2$; concentrations are indicated in the figure axes. Aliquots were resuspended in 50 mM Tris-buffered synthetic defined Zn-free media (SD-Zn), pH 7.4, and cell growth was measured by $OD_{600}$ after incubation for 48 h at 30 °C. **(B)** Data from the 1.6 µM Hist-5 row (red box in **A**) replotted as a bar graph to zoom in the nuanced modulatory effect of $Zn^{2+}$ under these conditions. Values and error bars represent the average and standard deviation from three separate biological replicates.

#### Figure S9. $Zn^{2+}$ addition blocks peptide internalization.

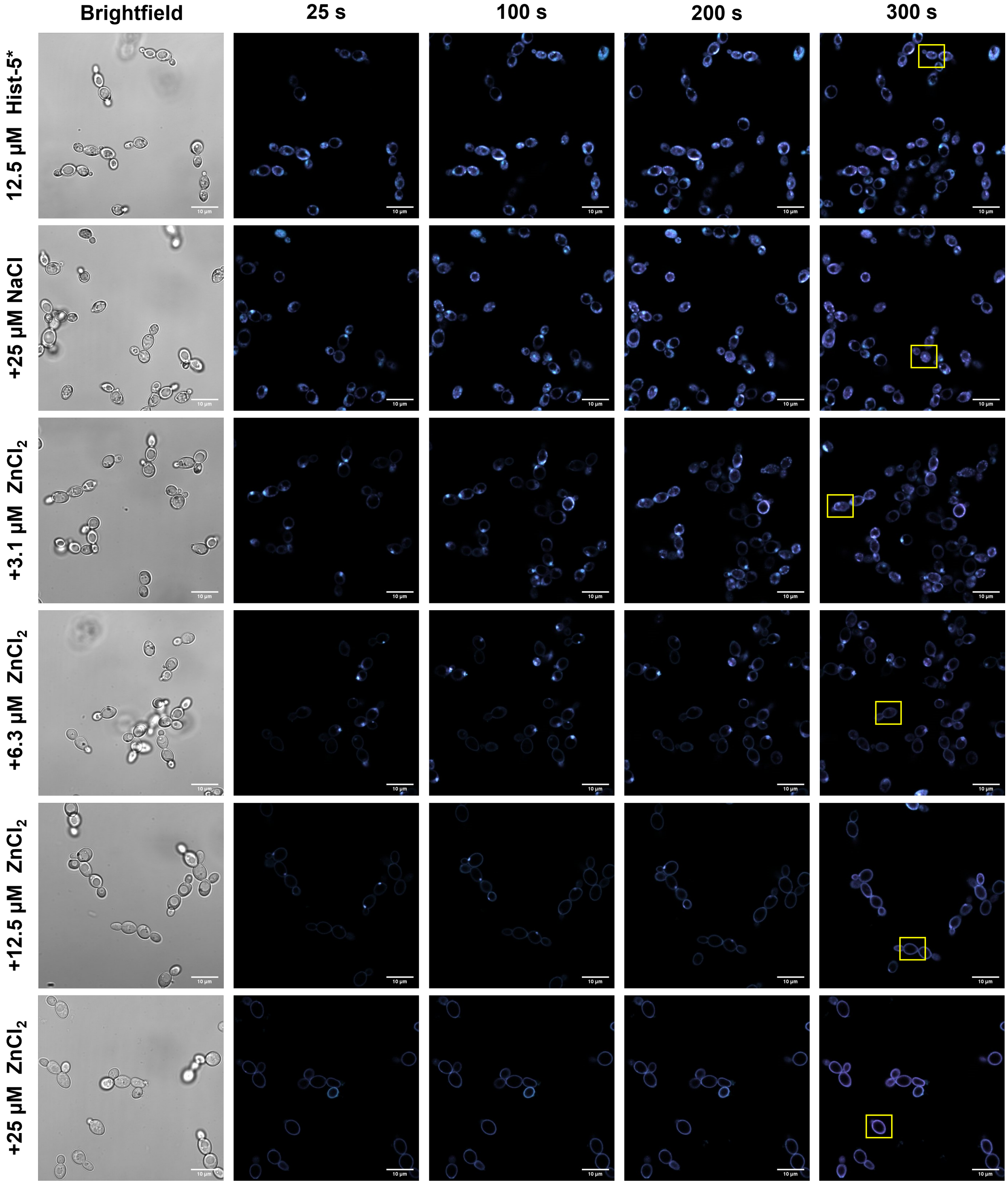


**Figure S9. $Zn^{2+}$ addition blocks peptide internalization.** Timelapse microscopy images of *C. albicans* cells treated with 12.5 µM Hist-5* alone, peptide+2 eq. NaCl, or varying concentrations of $ZnCl_2$ (0.25, 0.5, 1, or 2 eq. $Zn^{2+}$) at RT over 5 min in PPB.

**Figure S10. Zn-induced adherence of Hist-5* to the cell surface is long-lived.**

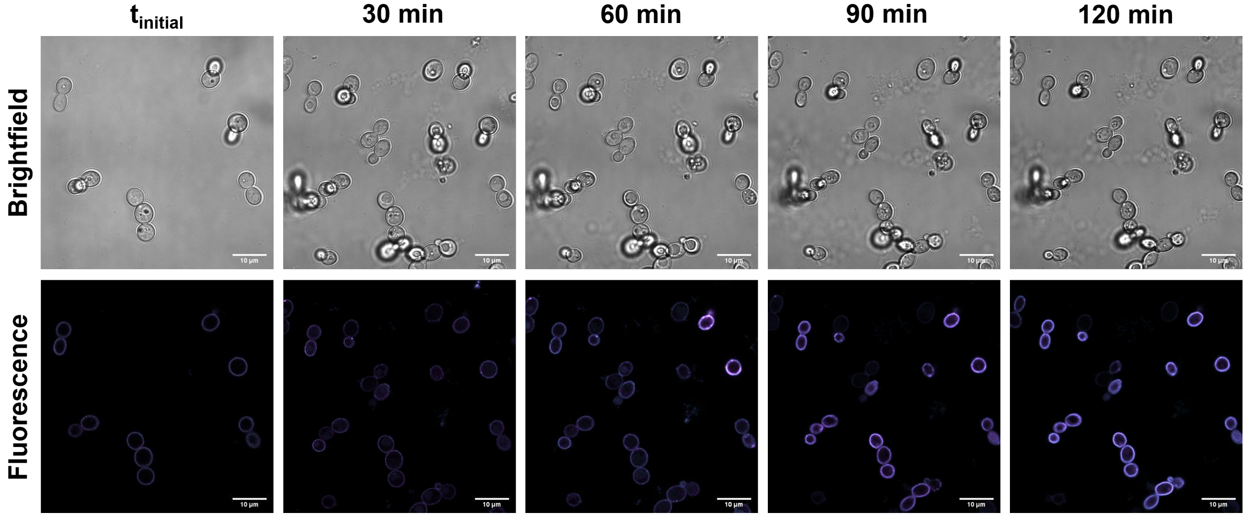


**Figure S10. Zn-induced adherence of Hist-5* to the cell surface is long-lived.** Timelapse microscopy images of *C. albicans* cells treated with 12.5 µM Hist-5* and 12.5 µM $ZnCl_2$ at RT over 2 h in PPB.

**Figure S11. Surface-bound effect of Hist-5* is Zn-specific.**

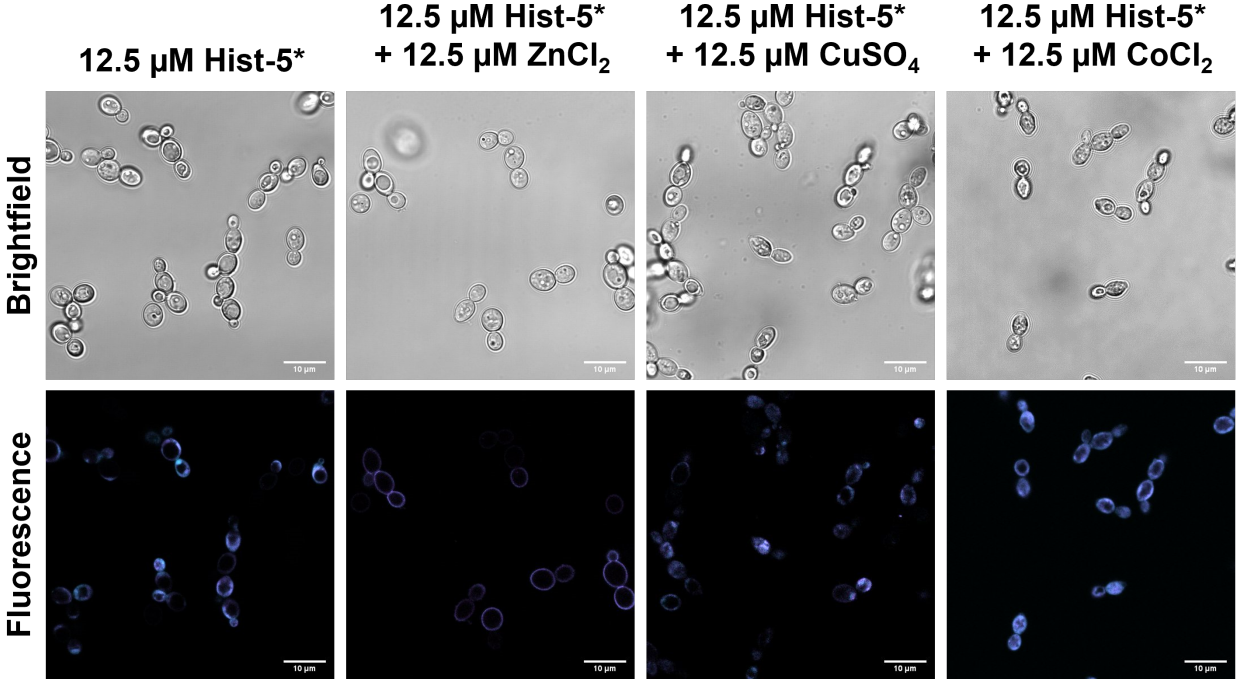


**Figure S11. Surface-bound effect of Hist-5* is Zn-specific.** Confocal microscopy images of *C. albicans* cells treated with 12.5 μM Hist-5* alone or 12.5 μM Hist-5*+12.5 μM $ZnCl_2$, 12.5 μM $CuSO_4$, or 12.5 μM $CoCl_2$ at RT for 5 min in PPB.

**Figure S12. In vitro formation of a ternary complex between Hist-5, ZQ, and $Zn^{2+}$.**

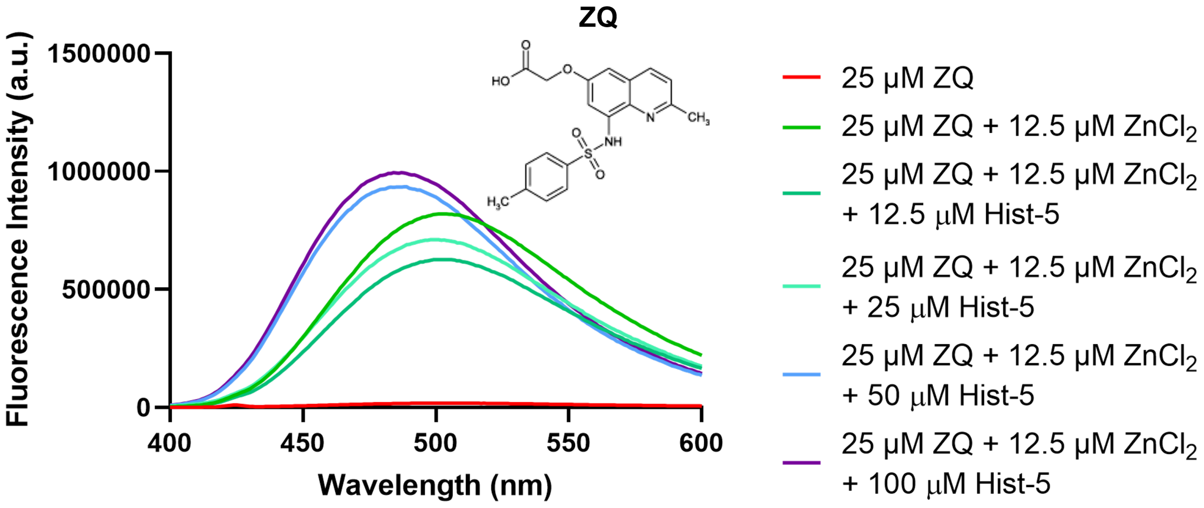


**Figure S12. In vitro formation of a ternary complex between Hist-5, ZQ, and $Zn^{2+}$.** Titration of Hist-5 into a solution containing 25 μM ZQ and 12.5 μM in $ZnCl_2$ PPB pH 7.4. Structure of ZQ free acid shown in the right-hand corner.

**Figure S13. EDTA and S100A12 protein do not induce membrane permeability.**

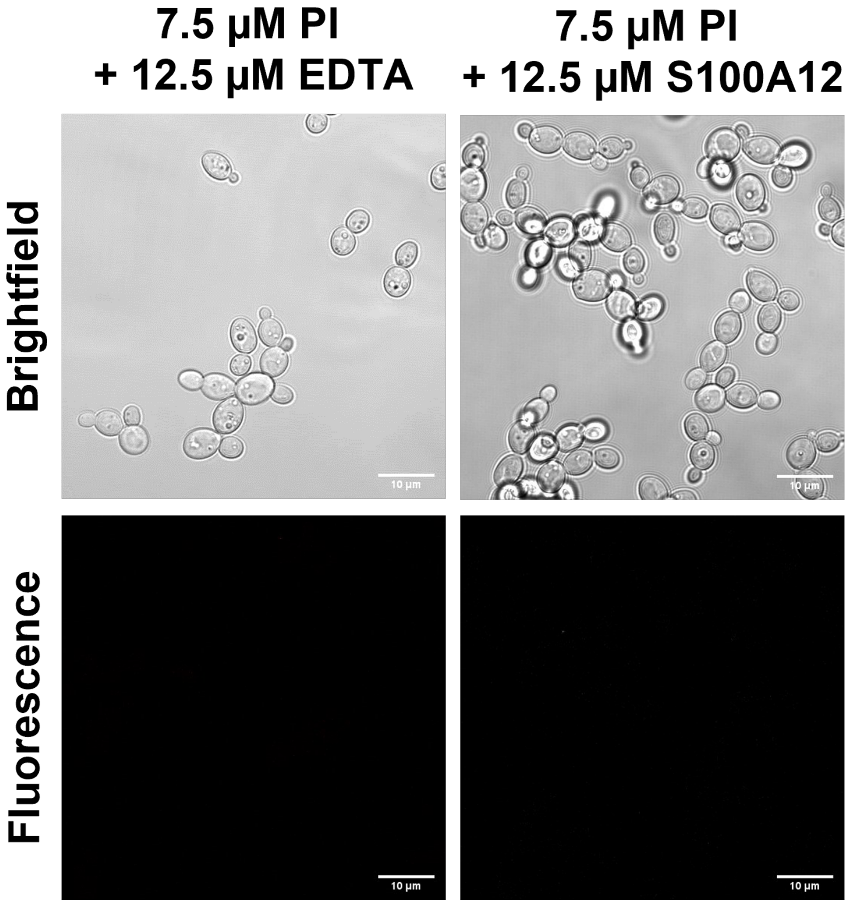


**Figure S13. EDTA and S100A12 protein do not induce membrane permeability.** Confocal microscopy images of *C. albicans* cells treated with 7.5 µM PI and 12.5 µM EDTA or 12.5 µM S100A12 at RT for 5 min in PPB.

**MATLAB script for image intensity analysis.** This script calculates the corrected total cell fluorescence (CTCF) value for each slice of a z-stack across all times within a confocal timelapse microscopy image.

$$CTCF = \frac{(Area\ of\ selected\ cell)*(Average\ pixel\ intensity\ within\ the\ area)}{(Area\ of\ background)*(Average\ pixel\ intensity\ of\ background)}$$

```
%% This script reads *.czi files containing image stacks for
% quantifying cell intensities.
% It uses open source bio-formats v5.3 dependencies for MATLAB
% extract bfmatlab.zip, downloadable from
% https://downloads.openmicroscopy.org/bio-formats/5.3.4/
% and copy file path below
% Written by Keerthi Anand
% 1/21/2020
% modified 3/22/20 to optimize run time
clc
close all
%%

%Paste full file path of extracted dependencies folder here
bfFolder='C:\Users\Joanna\Documents\MATLAB\bfmatlab';
addpath(genpath(bfFolder))

% Open dialog Box to load .czi file
[file,path,indx] = uigetfile('*.czi');

% Change to directory
cd(path)

% Extract all stacks and metadata
volume=bfOpen3DVolume([path file]);

%Extract data, which is located in the first cell
data=volume{1,1}{1,1};

%Number of stacks by 3 channels
maxStacks=size(data,3)/3;

% This is header info for data written out for each ROI/Cell
outputSummaryData={'Filename' 'Time' 'Frame' 'CH1 Area' 'CH1 Mean' 'CH1 ID' ...
    'CH1 CTFC' 'CH1 Mean Bgd' 'CH2 Area' 'CH2 Mean' 'CH2 ID' 'CH2 CTFC' ...
    'CH2 Mean Bgd'};

%Scale for area covered by each pixel, custom for your imaging field
factor=67.48^2/512^2;


% Second cell contains metadata, use this to find number of time points
% (numTotalTimePoints) and number z stacks (numTotalZStacks)
metadata = volume{1, 2};
metadataKeys = metadata.keySet().iterator();
```

```
numTotalTimePoints=1;
numTotalZStacks=1;

% Search through metadata to find fields containing number timepoints and
% z stacks
for z=1:metadata.size()
    key = metadataKeys.nextElement();
    value = metadata.get(key);

    if contains(key,'DimensionT')
        numTotalTimePoints=str2double(value);
    end
    if contains(key,'SizeZ')
        numTotalZStacks=str2double(value);
    end
    allMetdata{z,1}=key;
    allMetdata{z,2}=value;
    % fprintf('%s = %s\n', key, value)
end

% Specify colormaps for vizualizing channels
colormapMagenta=zeros([256 3]); % colormap for magenta
colormapMagenta(:,[1 3])=repmat(linspace(0,1,256)',[1 2]);
colormapCyan=zeros([256 3]); % colormap for cyan
colormapCyan(:,[2 3])=repmat(linspace(0,1,256)',[1 2]);

% Set intensity thresholds for chan 1 and 2 to get rid of noise
noiseThresholdChan1=15; % 15 works well but play around
noiseThresholdChan2=15;

% First run, set flag to 0
reuseROI=0;

% numTotalTimePoints = number  Timepoints
% numTotalZStacks = number z Stacks
% channels arranged tmpt1-z1-chans1,2,3;tmpt1-z2-chans1,2,3 ...
% Figure out how many images there are and iterate through them

% Allocate space for saving intensities through time
intensityThruTimeChan1=zeros(size(data,1),size(data,2),numTotalTimePoints);
intensityThruTimeChan2=zeros(size(data,1),size(data,2),numTotalTimePoints);

% idxDataEntry will be counter to add data to summary output file.
idxDataEntry=1;
for idxTime=1:numTotalTimePoints % Iterate through time

    %Allocate space
    intensityThruStacksChan1=zeros(size(data,1),size(data,2),numTotalZStacks);
    intensityThruStacksChan2=zeros(size(data,1),size(data,2),numTotalZStacks);

    % Iterate through z stacks
    for z=1:numTotalZStacks
        % Unique index to find data for particular
        % z-stack and time point
        imLoc=(idxTime-1)*numTotalZStacks*3+(z-1)*3;
```

```
        chan1=data(:,:,imLoc+1); % Mca fluorophore
        chan2=data(:,:,imLoc+2); % ABD fluorophore
        chan3=data(:,:,imLoc+3); % Bright field

        %Save intensity through stacks
        intensityThruStacksChan1(:,:,z)=chan1;
        intensityThruStacksChan2(:,:,z)=chan2;

        % Draw ROI for first time point and first z stack, we will
        % assume cells don't move too much
        if z==1 && idxTime==1 % if first z stack, draw ROIs
            figure(1)
            subplot(1,1,1)
            % Displays brightfield image
            imagesc(chan3)
            title(['File: ' file ' Bright Field'],'Interpreter', 'none')
            colormap(gca,'gray')
            axis image

            % First ROI is around cell of interest
            if reuseROI==0 || exist('main','var')==0
                roiCell = imfreehand();
                % Coordinates of ROI
                xyCell = roiCell.getPosition;
                hold on;
                % plot boundaries of ROI
                plot(xyCell(:, 1), xyCell(:, 2), 'b.', 'LineWidth', 2, 'MarkerSize',
10);
                mainCellMask = createMask(roiCell);
            end

            % Second ROI is around region in background
            if reuseROI==0 || exist('bgd1','var')==0
                roiBgd1 = imfreehand();
                % Coordinates of ROI
                xyBgd1 = roiBgd1.getPosition;
                hold on;
                % plot boundaries of ROI
                plot(xyBgd1(:, 1), xyBgd1(:, 2), 'r.', 'LineWidth', 2, 'MarkerSize',
10);
                bgdMask1 = createMask(roiBgd1);
            end

            % Third ROI is around region in background
            if reuseROI==0 || exist('bgd2','var')==0
                roiBgd2 = imfreehand();
                % Coordinates of ROI
                xyBgd2 = roiBgd2.getPosition;
                hold on;
                % plot boundaries of ROI
                plot(xyBgd2(:, 1), xyBgd2(:, 2), 'r.', 'LineWidth', 2, 'MarkerSize',
10);
                bgdMask2 = createMask(roiBgd2);

            end
```

```
        % Fourth ROI is around region in background
        if reuseROI==0 || exist('bgd3','var')==0
            roiBgd3 = imfreehand();
            % Coordinates of ROI
            xyBgd3 = roiBgd3.getPosition;
            hold on;
            % plot boundaries of ROI
            plot(xyBgd3(:, 1), xyBgd3(:, 2), 'r.', 'LineWidth', 2, 'MarkerSize',
10);
            bgdMask3 = createMask(roiBgd3);
        end

    else % if later zstack or tmpt, just calc values using existing ROI
        figure(1),

        % Display the three different channels as subplots
        h=subplot(2,2,1);
        imagesc(chan1,[0 50]),
        title('Channel 1')
        colormap(gca,colormapMagenta)
        axis image
        h2=subplot(2,2,2);
        imagesc(chan2,[0 50]),
        title({['T: ' num2str(idxTime) ' Z: ' num2str(z) ],'Channel 2'})
        colormap(gca,colormapCyan)
        axis image
        h3=subplot(2,2,3);
        imagesc(chan3),
        title('Bright Field')
        colormap(gca,'gray')
        axis image

        % Plot Cell ROI
        subplot(2,2,3)
        hold on
        plot(xyCell(:, 1), xyCell(:, 2), 'b.', 'LineWidth', 2, 'MarkerSize', 10);
        % Plot Background ROI 1
        hold on
        plot(xyBgd1(:, 1), xyBgd1(:, 2), 'r.', 'LineWidth', 2, 'MarkerSize', 10);
        % Plot Background ROI 2
        hold on
        plot(xyBgd2(:, 1), xyBgd2(:, 2), 'r.', 'LineWidth', 2, 'MarkerSize', 10);
        % Plot Background ROI 3
        hold on
        plot(xyBgd3(:, 1), xyBgd3(:, 2), 'r.', 'LineWidth', 2, 'MarkerSize', 10);
    end

    % Calculate the mean values of three background ROIs in chan 1 (Mca)
    meanNoiseIntensities(z,1,2)=mean(chan1(bgdMask1));
    meanNoiseIntensities(z,1,3)=mean(chan1(bgdMask2));
    meanNoiseIntensities(z,1,4)=mean(chan1(bgdMask3));
    meanNoiseIntensityBgdChan1=mean(squeeze(meanNoiseIntensities(z,1,2:end)));

    % Calculate the mean values of three background ROIs in chan 2 (ABD)
```

```
        meanNoiseIntensities(z,2,2)=mean(chan2(bgdMask1));
        meanNoiseIntensities(z,2,3)=mean(chan2(bgdMask2));
        meanNoiseIntensities(z,2,4)=mean(chan2(bgdMask3));
        meanNoiseIntensityBgdChan2=mean(squeeze(meanNoiseIntensities(z,2,2:end)));

        % Create a mask that calculates only the mean of data with
        % intensity above the background, and display this mask
        acceptMask=mainCellMask &
chan1>(meanNoiseIntensityBgdChan1+noiseThresholdChan1) &
chan2>(meanNoiseIntensityBgdChan2+noiseThresholdChan2);
        subplot(2,2,4)
        imagesc(acceptMask),
        title('Thresholded Mask')
        colormap(gca,'gray')
        axis image

        % Calulate avarge intensity of cell in chan 1
        meanNoiseIntensities(z,1,1)=mean(chan1(acceptMask));
        % Calulate avarge intensity of cell in chan 2
        meanNoiseIntensities(z,2,1)=mean(chan2(acceptMask));

        % Sum up area of cell
        areas=sum(sum(double(acceptMask)));

        % Compute Integrated Density chan 1
        ID(z,1)=factor*areas*meanNoiseIntensities(z,1,1);
        % Compute Integrated Density chan 1
        ID(z,2)=factor*areas*meanNoiseIntensities(z,2,1);

        % Compute Corrected Total Cell Fluorescence chan 1
        CFTC(z,1)=factor*areas*(meanNoiseIntensities(z,1,1)-
meanNoiseIntensityBgdChan1);
        % Compute Corrected Total Cell Fluorescence chan 2
        CFTC(z,2)=factor*areas*(meanNoiseIntensities(z,2,1)-
meanNoiseIntensityBgdChan2);

        % Uncomment for display of  ID and CFTC values (may slow down script)
        % figure(1)
        % subplot(2,2,1)
        % xlabel({['ID: ' num2str(ID(i,1))],['CFTC: ' num2str(CFTC(i,1))]})
        % subplot(2,2,2)
        % xlabel({['ID: ' num2str(ID(i,2))],['CFTC: ' num2str(CFTC(i,2))]})

        % update figure
        drawnow

        % Save summary data to output file and increment counter
        idxDataEntry=idxDataEntry+1;
        outputSummaryData(idxDataEntry,:)={file idxTime z areas*factor
meanNoiseIntensities(z,1,1) ID(z,1) CFTC(z,1) meanNoiseIntensityBgdChan1 areas*factor
meanNoiseIntensities(z,2,1) ID(z,2) CFTC(z,2) meanNoiseIntensityBgdChan2};
    end
    % Save Intensity through stacks
    intensityThruTimeChan1(:,:,idxTime)=sum(intensityThruStacksChan1,3);
    intensityThruTimeChan2(:,:,idxTime)=sum(intensityThruStacksChan2,3);
```

```
end

%% Saves ROI and stat data under custom suffix
time=clock;
time=num2str(time(1:end-1));
time(time == ' ') = [];

% Prompt for user suffix
prompt = {'Enter ROI identifier eg Cell1 or C1'};
dlgTitle = 'Input';
dims = [1 40];
defInput = {[time '_C1'],'hsv'};
answer = inputdlg(prompt,dlgTitle,dims,defInput);

fileSuffix=[answer{1}];
saveFilename=[file(1:end-4) '_' fileSuffix '.mat'];

% Save data to .mat file, load this to reuse masks, change reuse flag to 1
save(saveFilename,'mainCellMask','bgdMask1','bgdMask2','bgdMask3','outputSummaryData'
,'intensityThruTimeChan1','intensityThruTimeChan2','noiseThresholdChan1','noiseThresh
oldChan2','acceptMask','xyCell','xyBgd1','xyBgd2','xyBgd3')
```
